# Ontogenetic consequences of developmental temperature in amphibians: simultaneous gains in heat tolerance and cumulative costs to stress physiology

**DOI:** 10.64898/2026.03.01.708817

**Authors:** Ingrid R. Miguel, Pablo Burraco, Carlotta Hakemann, Lara Keunecke, Colette Martin, Natasha Kruger, Katharina Ruthsatz

**Affiliations:** Institue for Cell– and Neurobiology, Technische Universität Braunschweig, Mendelssohnstraße 4, 38106 Braunschweig, Germany; Zoology Department, Biology Institute in Federal University of Rio de Janeiro, Centro de Ciências e Saúde (CCS) – Avenida Brigadeiro Trompowski, s/n°, Cidade Universitária – Ilha do Fundão CEP: 21941-902 – Rio de Janeiro, RJ, Brazil; Vertebrate Department, Herpetology Section, National Museum in Rio de Janeiro, Brazil; Doñana Biological Station (CSIC), 41092 Sevilla, Spain; Animal Behaviour and Wildlife Conservation Group, School of Life Sciences, University of Wolverhampton, Wolverhampton WV1 1LY, United Kingdom

**Author notes:** N. Kruger and K. Ruthsatz share the last authorship due to equal contribution. Corresponding author: **Katharina Ruthsatz**;. Current address: Ecology, Evolution and Development Group, Department of Wetland Ecology, Doñana Biological Station (CSIC), 41092 Sevilla, Spain.

**Keywords:** global warming, heat stress, early life stage effects, trade-off, metabolism, stress physiology, thermal acclimation, thermal tolerance

## Abstract

Heat tolerance is critical for ectotherms facing environmental temperature variability, yet how it varies across life stages, and whether trade-offs occur between temperature-induced developmental plasticity and heat tolerance, remain unclear, particularly in organisms undergoing metamorphosis which represent 95% of all animal species. We examined how early-life thermal conditions shape growth, development, survival, acclimation capacity, heat tolerance, and energy allocation across ontogeny in the African clawed frog (*Xenopus laevis*), reared at six constant temperatures (17–32°C). We tested whether higher developmental temperatures generate trade-offs between accelerated growth and heat tolerance, and the consequences for post-metamorphic resilience to extreme heat. Rearing at 32°C was lethal before metamorphosis. At non-lethal warm temperatures (17–29°C), larvae and juveniles simultaneously accelerated development, maintained growth, and enhanced heat tolerance. However, juveniles reared at 29°C showed reduced survival, elevated corticosterone responses to acute stress, and diminished acclimation capacity, indicating increased energetic demands and constrained metabolic flexibility. These findings show that amphibians can integrate developmental plasticity with plastic adjustments in heat tolerance, but that such strategies incur cumulative physiological costs. By adopting an across-life-stage approach, our study highlights energy-allocation constraints that may limit population persistence under climate warming in species with complex life cycles.

## 1. Introduction

Climate change is altering environmental conditions at an unprecedented rate, as rising mean temperatures and more frequent heat waves pose major challenges to wildlife (IPCC, 2023). Ectothermic organisms are particularly vulnerable to these changes (Paaijmans et al., 2013), as their physiological processes are directly influenced by external temperatures due to their limited ability to regulate internal body temperature (Hochachka & Somero, 2002). In species with complex life cycles such as those undergoing metamorphosis, thermal conditions experienced during development can produce carry-over effects from one stage to the next (Kingsolver et al., 2011), as well as cumulative effects that build across multiple stages, influencing growth, physiological performance, and survival throughout ontogeny (Monaghan, 2008). While such effects can extend beyond individuals (Burton & Metcalfe, 2014), how thermal environments cumulatively experienced across early life stages influence resilience to acute stress later in life remains poorly understood (Metcalfe & Monaghan, 2001). Specifically, little is known about the influence of developmental temperatures on an organism’s sensitivity to or ability to cope with acute thermal fluctuations later in life, a key factor in determining resilience to climate change (Nord & Giroud, 2020). Previous research has shown that developmental thermal environments can influence post-metamorphic growth, morphology, and survival (e.g., Gomez-Mestre et al., 2010; Ruthsatz et al., 2020a; Pottier et al., 2022a; Ohmer et al., 2023). However, comparatively less is known about how these effects unfold across multiple life stages and whether early-life thermal plasticity generates cumulative costs for later acute stress resilience.Addressing this knowledge gap is needed for advancing our predictions of populations and species responses to climate change, ultimately supporting more effective conservation and management strategies.

Thermal tolerance is a key physiological trait that defines the range of temperatures over which organisms can function (Huey & Stevenson, 1979; Angilletta et al., 2002). Thermal tolerance is often quantified with thermal performance curves, which illustrate how traits such as metabolism, locomotion, and growth vary with temperature (Sinclair et al., 2016). These curves peak at an optimum and drop sharply beyond critical limits, reaching the critical thermal maximum (CT_max_), the upper temperature at which survival is no longer possible (Angilletta, 2009). In this context, CT_max_ represents an organism’s heat tolerance, and is therefore an important component of vulnerability to rising temperatures and more frequent extreme heat events (Pörtner, 2001; Sinclair et al., 2016). However, species and population persistence are not determined by thermal tolerance alone, but also depend on additional factors such as thermal fertility limits, behavioral heat avoidance, life-history responses, and demographic recovery following thermal disturbances. Similarly, absolute heat tolerance alone may not be sufficient to buffer organisms against rapid environmental change. Instead, an individual’s ability to adjust its heat tolerance through thermal acclimation or acclimatization in response to changing thermal conditions plays a crucial role in thermal resilience (Calosi et al., 2008; Seebacher et al., 2015; but see: Gunderson & Stillman, 2015). This acclimation capacity may allow organisms to extend their functional thermal range, providing an important buffer against rising temperatures (Seebacher et al., 2015). However, the extent of acclimation varies widely among species and populations and is influenced by genetic, ecological, and developmental factors (Sunday et al., 2011; Rohr et al., 2018; Pinsky et al., 2022; Ruthsatz et al., 2024). Importantly, thermal tolerance is not constant across an organism’s life cycle (Dahlke et al., 2020; Ruthsatz et al., 2022a), and certain stages may exhibit weaker acclimation capacity (Pottier et al., 2022a; Weaving et al., 2022; Ruthsatz et al., 2024). The life stage with the lowest heat tolerance may act as a thermal bottleneck, constraining population persistence under climate warming (Dahlke et al., 2020; Ruthsatz et al., 2022a; but see: Pottier et al., 2022b). This risk is greatest when heat tolerance is close to current environmental temperatures, narrowing thermal safety margins and leaving organisms more vulnerable to acute heat stress (Gunderson & Stillman, 2015). Identifying thermally vulnerable life stages and the extent to which early-life thermal conditions shape long-term thermal resilience, is essential for predicting species’ responses to climate change.

Beyond acclimation capacity in heat tolerance, plasticity in growth and development represents a key and often adaptive trait of species with complex life cycles, allowing developmental trajectories to be shaped by environmental challenges, regardless of potential long-term consequences (Wilbur, 1980; Denver & Middlemis-Maher, 2010). Such developmental plasticity in both physiological and metamorphic traits is highly adaptive in temporary or rapidly changing habitats, where individuals must adjust their development and thermal performance to optimize survival and fitness (Newman, 1992; Rudolf & Rödel, 2007; Chevin & Hoffmann, 2017). However, this raises an important question: *Can individuals simultaneously invest in rapid growth and development while also maintaining high heat tolerance, or does energy allocation impose an unavoidable trade-off between these two responses?* Maintaining a high heat tolerance is energetically costly, as it requires upregulation of protective cellular mechanisms and increased metabolic activity (Sokolova, 2018; Chung & Schulte, 2020; Sokolova, 2023), which could divert resources away from growth and development. If individuals prioritize rapid development and early metamorphosis, they may allocate fewer resources toward developing thermal resilience, here referring to the capacity to tolerate and recover from acute heat stress, potentially compromising their ability to withstand acute heat stress later in life (Sokolova, 2013). Conversely, if energy is primarily directed toward building heat tolerance, growth may slow (Nord & Giroud, 2020), leading to delayed metamorphosis or smaller size at emergence, both of which could reduce survival and fitness in later life stages (Werner, 1986; Rowe & Ludwig, 1991; Blaustein et al. 2010). These trade-offs are particularly relevant in species undergoing metamorphosis such as amphibians, a period of extensive physiological restructuring and high metabolic demand (Ruthsatz et al., 2019), which may further constrain the ability to allocate energy towards both growth and thermal resilience. Beyond these immediate trade-offs, carry-over effects from early-life thermal conditions can influence post-metamorphic performance and survival (Monaghan, 2008). If early thermal stress leads to energy limitations at metamorphosis, juveniles may face reduced stress resilience, lower thermal performance, or diminished survival probability (Scott et al. 2007). These effects could scale up to impact population dynamics, as lower juvenile survival rates and impaired reproductive success may reduce population persistence, particularly in species already facing climate-induced habitat loss. Therefore, understanding whether and how organisms balance developmental plasticity in growth with thermal resilience across life stages is crucial for predicting their long-term responses to climate change.

In this study, we investigated how thermal conditions experienced during early life stages (before, during, and after metamorphosis) shape amphibian growth, development, thermal physiology, and survival. Using the African clawed frog (*Xenopus laevis*) as a model, we experimentally manipulated rearing temperatures to test their effects on larval developmental timing and growth, as well as energy allocation and thermal tolerance from larvae to post-metamorphic juveniles. We specifically addressed whether early-life thermal environments generate trade-offs between rapid growth and heat tolerance, and whether cumulative effects influence juvenile resilience to acute thermal stress. We hypothesized that higher developmental temperatures would accelerate growth and metamorphosis but constrain energy available for thermal protection, thereby reducing overall survival and juvenile stress resilience. To assess this, we used corticosterone (CORT) as a physiological indicator of acute stress responsiveness, predicting that juveniles would exhibit stronger CORT responses and lower survival. By adopting an across-life-stage approach, our study fills a critical gap in understanding the ontogeny of thermal resilience and the energy-allocation trade-offs that may limit population persistence under climate change, a dimension rarely explored in species with complex life cycles.

## 2. Materials and Methods

Full methodological details, definitions of physiological terms, and a detailed overview of the parameters measured at each life stage with corresponding sample sizes (Figure S1), are provided in the Supporting Information.

### 2.1 Animal husbandry and experimental design

We obtained three clutches (i.e., full-sib families) of the African clawed frog (*Xenopus laevis*) from the captive breeding facility of the Universitätsklinikum Hamburg Eppendorf (UKE, Martinistr. 52, 20246 Hamburg, Germany) in February 2024. Until the synchronous hatching and developmental stage NF45 (i.e., onset of exotrophic feeding), each clutch was kept separately at 22 °C in dechlorinated water, matching the long-term rearing temperature of the parental stock.

A three-phase experimental design was chosen to assess the effects of acclimation temperature during ontogeny on larvae (phase 1; developmental stage NF57, *i.e.*, all five toes separated, Nieuwkoop et al., 1994), metamorphs (phase 2; developmental stage NF66, *i.e.,* metamorphosis is completed, tail fully resorbed, Nieuwkoop et al., 1994), and juveniles (phase 3; 150 days after hatching). The experiment was conducted in one climate chamber (Kälte-Klimatechnik-Frauenstein GmbH, Germany) with a 14:10 h light:dark cycle and a mean (±SD) air temperature of 17 (± 0.5) °C. Our experimental setup had six different treatments consisting in constant temperature (i.e., mean±SD) of 17.0±0.5, 20.0±0.2, 23.0±0.2, 26.0±0.1, 29.0±0.1, and 32.0±0.2 °C. Each treatment was replicated six times, for a total of 36 aquaria. The selected rearing temperatures fall within the broad thermal range experienced by *X. laevis* tadpoles in their native range, where tadpoles show a broad 80% thermal performance breadth from 13.1°C to 36.7°C (Ginal et al., 2023). This allowed us to test how different, yet biologically relevant, early-life thermal environments influence subsequent development and thermal resilience. Across all life stages, water was changed twice a week with aerated aged tap water pre-heated to the treatment-specific temperature. Dead animals were removed from the aquaria. The experiments ran for 23 weeks.

*Phase 1: larvae* – Five larvae from each clutch were randomly allocated to each of the 36 standard 12-L aquaria, filled with 9 L of de-chlorinated water. This design ensured that all three clutches were equally represented in each aquarium and treatment group. Each aquarium housed 15 larvae (15 larvae × 36 aquaria = 540 larvae in total; larval density: 1.66 larvae × L^-1^). This density was selected to maintain larvae below densities commonly associated with crowding effects, while allowing group rearing, which is appropriate for *X. laevis* tadpoles because they naturally aggregate and show schooling behavior (Katz et al., 1981; Lopez et al., 2021). For each of the six experimental temperatures, six aquaria were maintained using independent adjustable heating elements (JBL PROTEMP S 25, 25 W, JBL GmbH & Co. KG, Germany). Each aquarium was equipped with an air stone connected to a pump to ensure aeration of the holding water and a proper distribution of the food particles. Larvae were fed a mix of 50 % high-protein powdered fish food (Sera micron, Sera, Germany) and 50 % Spirulina algae powder *ad libitum*. When larvae reached developmental stage NF57, a subset of larvae from each aquarium was randomly selected for heat tolerance (N=5 individuals per aquarium) and metabolism measurements (N=3 individuals per aquarium). All surviving animals remained in their aquaria until the completion of metamorphosis (developmental stage NF66). At 32 °C, no larvae survived to the first sampling stage; therefore, all measurements are based on animals reared at 17, 20, 23, 26, and 29 °C.

*Phase 2: metamorphs* – When approximately 50% of individuals in an aquarium had completed metamorphosis (developmental stage NF66), three metamorphosed individuals were randomly selected for heat-tolerance measurements. The remaining animals were kept in their larval groups in accordance with animal welfare guidelines. After completing metamorphosis, all remaining animals were transferred to plastic boxes (Surplus Systems Eurobox, 6000 mm x 4000 mm x 2200 mm) equipped with a minimum of one and a maximum of three adjustable heating elements (JBL PROTEMP S 25, 25 W, JBL GmbH & Co. KG, Germany) to maintain the respective acclimation temperature and an air stone for constant aeration of the holding water. Metamorphs from all six larval aquaria per temperature treatment were pooled in two to three boxes and kept at a density of 1.66 animals × L^-1^. Each box was equipped with clay-made shelters for enrichment (Archard et al., 2012) and covered with a plastic mesh. Animals were fed *ad libitum* with living bloodworms (*Chironomus aprilinus*) as well as high-protein frog food granules (Tetra ReptoFrog Granules, Tetra GmbH, Germany) and were kept in the boxes until the final measurements.

*Phase 3*: *juveniles* – Final measurements were conducted at 150 days after hatching. This sampling point was chosen in order to have a comparable age and time period for growth from hatching for all animals across all temperature treatments. From all surviving animals (N=133), a minimum of five (due to survival limitations) and a maximum of 16 juveniles per temperature treatment were randomly selected for heat stress resilience assessments (see survival results below). All remaining juveniles were assigned to heat tolerance measurements (N=73).

### 2.2 Life history variables

Ontogenetic stage was determined using morphological criteria following Nieuwkoop et al., (1994). Age was expressed as days after hatching (dah). Developmental rate was calculated as the number of Nieuwkoop and Faber (NF) stages progressed per day from NF45 (start of experiment, 6 dah) to NF57 (larval stage). Snout–vent length (SVL), and survival body mass were recorded at NF45, NF57, NF66 (metamorphs), and at 150 dah (juveniles). Growth rates (mg d ¹) were calculated as the change in body mass between consecutive stages, divided by the elapsed time. Because developmental temperature influenced the timing of metamorphosis, juveniles differed in the duration of the post-metamorphic period at this standardized sampling point. To account for this, juvenile growth rate was calculated relative to the time elapsed since metamorphosis.

### 2.3 Heat tolerance assessments

We measured critical thermal maximum (CT_max_) as a proxy for heat tolerance at three life stages, late larvae (NF57), metamorphs (NF66), and juveniles (150 dah), following Sinai et al., (2024). Briefly, CT_max_ was determined by increasing water temperature from the acclimation temperature at +0.1 °C min ¹ until loss of locomotor function (i.e., loss of rightening response). After testing, animals recovered at intermediate temperatures before being euthanized, measured, photographed, and preserved for later fat body dissection.

#### 2.3.1 Life stage-specific acclimation capacity in heat tolerance

Acclimation capacity in CT_max_ was estimated within each life stage using the acclimation response ratio (ARR) (Claussen, 1977):

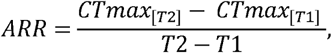

where T1 and T2 are the lowest (17 °C) and highest (29 °C) acclimation temperatures, respectively. Higher ARR values indicate greater thermal plasticity; an ARR of 1.0 corresponds to full acclimation (Morley et al., 2019).

### 2.4 Metabolism measurements

We measured oxygen consumption in three randomly selected larvae (NF57) per aquarium from all surviving temperature treatments (N = 90; 3 larvae × 30 aquaria)using closed respirometry to estimate routine metabolic rate (RMR) during the natural activity phase (08:30–22:30 h; Ruthsatz et al., 2019). Individuals were placed in sealed 30 mL glass vials filled with aerated, filtered water and equipped with optical oxygen sensors connected to a multi-channel oxygen measurement system (Oxy-4 SMA; PreSens Precision Sensing GmbH, Germany). Oxygen concentration was recorded every 15 s for 30 min, beginning 10 min after transfer to the chamber. Trials were run at the acclimation temperature and consecutively repeated at +2 °C and +4 °C above acclimation temperature to assess acute thermal sensitivity (Sinai et al., 2024). Thereafter, the dry blotted body mass of each specimen was rapidly determined before the larvae were returned to their aquaria. All larvae survived the measurements (i.e., no mortality after 24 h). Full details on vial preparation, sensor calibration, background respiration correction, oxygen depletion thresholds, and temperature-ramping procedures are provided in the Supporting Information.

#### 2.4.1 Routine metabolic rate calculations

For each individual, RMR was calculated as the slope of the linear least-squares regression of dissolved oxygen decline over 30 min, excluding activity peaks and the first 5 min to reduce handling effects (Hastings & Burggren, 1995). Rates were corrected for animal volume (Giacomin et al., 2019) and expressed as ml O□ h□^1^ g□^1^ dry-blotted body mass. Analyses were performed using PreSens Oxygen Calculator Software (PreSens Precision Sensing GmbH) and following Ruthsatz et al., (2022b).

#### 2.4.2 Thermal sensitivity of RMR

Thermal sensitivity was expressed as Q□□, calculated from respiration rates between successive temperature steps, with higher Q□□ values indicating greater increases in RMR at elevated temperatures. Mean Q_10_ values were calculated from the Q_10_ values between the three temperature steps (Sinai et al., 2024).

### 2.5 Acute heat stress experiment

Juveniles at 150 dah selected for heat stress resilience assessments were randomly assigned to a acute heat stress (N=4-8/temperature treatment) or control group (N=2-8/temperature treatment). Animals were kept individually in plastic boxes filled with 1.66 L of tap water pre-heated to the treatment-specific temperature. A total of 60 boxes were equipped with an adjustable heating element as well as an air stone and covered with a plastic mesh.

To impose an acute extreme-temperature challenge, we increased the temperature at a rate of 0.5°C per hour to 5°C above the original acclimation temperature. Animals were exposed to this acute heat stress treatment or control conditions for six days, following a previously published amphibian heat-stress protocol (Ujszegi et al., 2022).During the acute heat stress, we changed the water in the boxes every other day with aerated aged tap water pre-heated to the treatment-specific temperature and fed animals *ad libitum* with bloodworms and frog food granules. Control individuals experienced the same handling and treatment conditions. At the end of the six-day long heat wave period, we determined baseline waterborne CORT release rate at respective treatment temperature and body mass of each specimen. Eleven animals died during the experiment, including three control individuals and eight individuals exposed to acute heat stress.

### 2.6 CORT release, extraction, and quantification

We quantified waterborne CORT release at the end of the heat stress resilience assessments using the established protocol by Gabor et al., (2013). Each animal was placed for 1 h in pre-heated, aerated water at its treatment temperature, after which the water sample was collected and stored at −20 °C until processing. CORT was extracted via C18 solid-phase extraction and quantified using a DetectX Corticosterone ELISA kit (K014-H5, Ann Arbor, MI, USA), following manufacturer instructions.

In total, we ran one plate with 38 samples (i.e., 37 animal samples, 1 pooled control). Hormonal concentration of samples was measured in duplicate, and values were calculated from standard curves based on calibration standards provided with the DetectX kit using MyAssays online tools. The mean coefficient of variation of duplicates for all samples was 6.61 %. Average R^2^ for the 4PLC fitting curve was 0.999. CORT release rates were expressed as pg × g□^1^ × h□^1^ by standardizing concentrations to individual body mass (Gabor et al., 2013).

### 2.7 Life-stage specific fat reserves

The major triglyceride storage in amphibians is located in the abdominal area forming fat bodies, which largely explain metamorphic success and post-metamorphic survival (Scott et al., 2012; Wright et al., 2011). To assess life stage-specific effects of temperature on fat reserves, frozen specimens were thawed, blotted, and weighed on a high-precision balance (Ohaus VP-114CN Voyager Analytical Balance, Spain). Fat bodies were then dissected, blotted, and weighed on the same balance to the nearest 0.1 mg. Relative fat body size was calculated from individual mass of fat body (mg) divided by individual body mass (mg) and expressed as mg × mg body mass^−1^ (Ruthsatz et al., 2020b).

### 2.8 Statistical analysis

All models were fitted in R (R Core Team) using *lmer* from the lme4 package (Bates et al., 2015) for Gaussian linear mixed models, *glmer* (lme4) for binomial generalized linear mixed models, and *logistf* from the logistf package (Heinze et al., 2024) for Firth’s penalized logistic regression, applied only where noted below. Tank (or Replicate, for the survival dataset) was included as a random intercept in all mixed models. Temperature was treated as an ordered categorical factor rather than a continuous predictor or an unordered factor, since rearing temperature was implemented as a discrete set of experimental treatments. Model assumptions were assessed using simulation-based residual diagnostics (DHARMa; Hartig, 2022), including tests for uniformity of scaled residuals and overdispersion.

For each response variable, we fitted a single, biologically motivated model specified a priori. Significance of each fixed effect was assessed directly using Type II (or, for models including an interaction term, Type III) Wald chi-square tests (car::Anova; Fox & Weisberg, 2019). Post hoc pairwise comparisons across temperature treatments were performed using Tukey-adjusted tests via emmeans (Lenth, 2023), with the exception noted below for survival to 150 days after hatching.

A sixth treatment (32°C) was included in the original experimental design but resulted in complete mortality no individuals from this treatment survived to be included in any subsequent measurement or analysis. All models reported here therefore span the five temperatures at which data were available (17-29°C).

*Life-history traits.* Snout-vent length, body mass, and growth rate were analyzed separately for larvae, metamorphs, and juveniles, with temperature (ordered factor) as the independent variable. We additionally tested tank-level survival rate (a proxy for final density) and developmental age as covariates for each trait. Developmental age (dah) was consistently and strongly collinear with temperature across all traits and life stages tested (Pearson r = 0.88– 0.97), reflecting that temperature is the primary driver of developmental timing in this design; age was therefore excluded from all final models to avoid destabilizing the temperature effect. Tank-level survival rate showed no meaningful collinearity with temperature (VIF < 2) and was retained as a covariate throughout.

Rather than modeling developmental rate directly, we modeled days to reach the target developmental stage (NF57 for larvae), since developmental rate is a deterministic transformation of this count (rate = stages progressed ÷ days) and, because every individual progressed through the same fixed number of stages, produced a small number of discrete, clustered values that were poorly suited to standard continuous-response families (confirmed via residual diagnostics under Gaussian, Gamma, and Beta families). Days-to-stage was modeled as a Gaussian linear mixed model with temperature as the fixed effect; estimated marginal means and pairwise comparisons are additionally reported on the developmental rate scale (back-transformed as rate = 12/days) for biological interpretability, though inference (standard errors, test statistics, p-values) was performed on the days-to-stage scale on which the model was fit.

*Heat tolerance (CTmax).* CTmax was analyzed using a linear mixed model including life stage, temperature, their interaction, and body mass as a covariate..

*Metabolic traits*. Mean Q□□ and routine metabolic rate (RMR) were measured in a subsample of larvae and modeled with temperature as the sole fixed effect.

*Relative fat body mass.* Relative fat body mass was modeled as a function of life stage, temperature, their interaction, and growth rate (a combined measure spanning larval, metamorph, and juvenile growth) as a covariate. No juveniles survived to be sampled for this measurement at 29°C, so this model was fit across the temperatures at which all three life stages were represented.

*Corticosterone (CORT) release rate*. We first tested whether baseline (control) and post-heatwave CORT release rate were each predicted by rearing temperature, fitting these as two separate linear mixed models (one restricted to control individuals, one to post-heatwave individuals). To evaluate the acute heatwave effect specifically, we additionally fitted separate linear mixed models within each rearing temperature, with heatwave exposure (control vs. heat-challenged) as the fixed effect.

*Survival.* Survival to NF57 (larval stage) and NF66 (metamorph stage) was analyzed using binomial mixed-effects logistic regression, with the count of survivors and deaths as the response and tank/replicate as a random intercept; both models converged without evidence of separation. Survival to 150 days after hatching showed complete separation (100% survival in one temperature treatment), preventing reliable estimation via the standard mixed-effects approach; the random intercept for tank/replicate in that model was estimated at zero variance, indicating no random-effect information was lost by its omission, so we instead used Firth’s penalized logistic regression without a random effect for this endpoint. Because emmeans does not support Firth models, pairwise comparisons for this endpoint were obtained by fitting separate two-group models for each pair of temperatures; these p-values are not jointly Tukey-adjusted, unlike all other pairwise comparisons reported. Separately, all individuals reared at 32°C died at the very start of the experiment, before reaching any survival checkpoint; this temperature is therefore not represented in any of the survival models and is instead reported as a discrete early-mortality finding.

## 3. Results

Full model results, including model structure, test statistics, degrees of freedom, and p-values for each response variable, are reported in Table S1, while detailed Tukey-adjusted post hoc comparisons are provided in Table S2 (Firth-based comparisons for survival to 150 days after hatching, which used unadjusted p-values, are reported separately in Table S3).

### 3.1. Life-history traits

*Larvae.* Temperature significantly influenced all larval life-history traits (Table S1). Days to reach NF57 decreased progressively with temperature (χ²=669.17, df=4, p<0.001); all pairwise comparisons for larval developmental rate among 17, 20, and 23°C were significant (p<0.001), as were comparisons involving 26°C and 29°C (p<0.001–0.015), with development slowest at 17°C and fastest at 26–29°C. SVL and body mass both decreased significantly with temperature (SVL: χ²=79.33, df=4, p<0.001; mass: χ²=251.51, df=4, p<0.001): individuals at 17°C and 20°C were both significantly larger and heavier than individuals at 23, 26, and 29°C (all pairwise p<0.001–0.002). Larval growth rate was significantly affected by temperature overall (χ²=9.81, df=4, p=0.044), although no individual pairwise comparison between temperatures remained significant after Tukey correction (Figure 1). Tank-level survival rate (a proxy for density) additionally predicted larval growth rate independent of temperature (χ²=5.26, df=1, p=0.022), with growth rate increasing as tank-level survival rate increased (β=0.462, SE=0.201).

**Figure 1.**
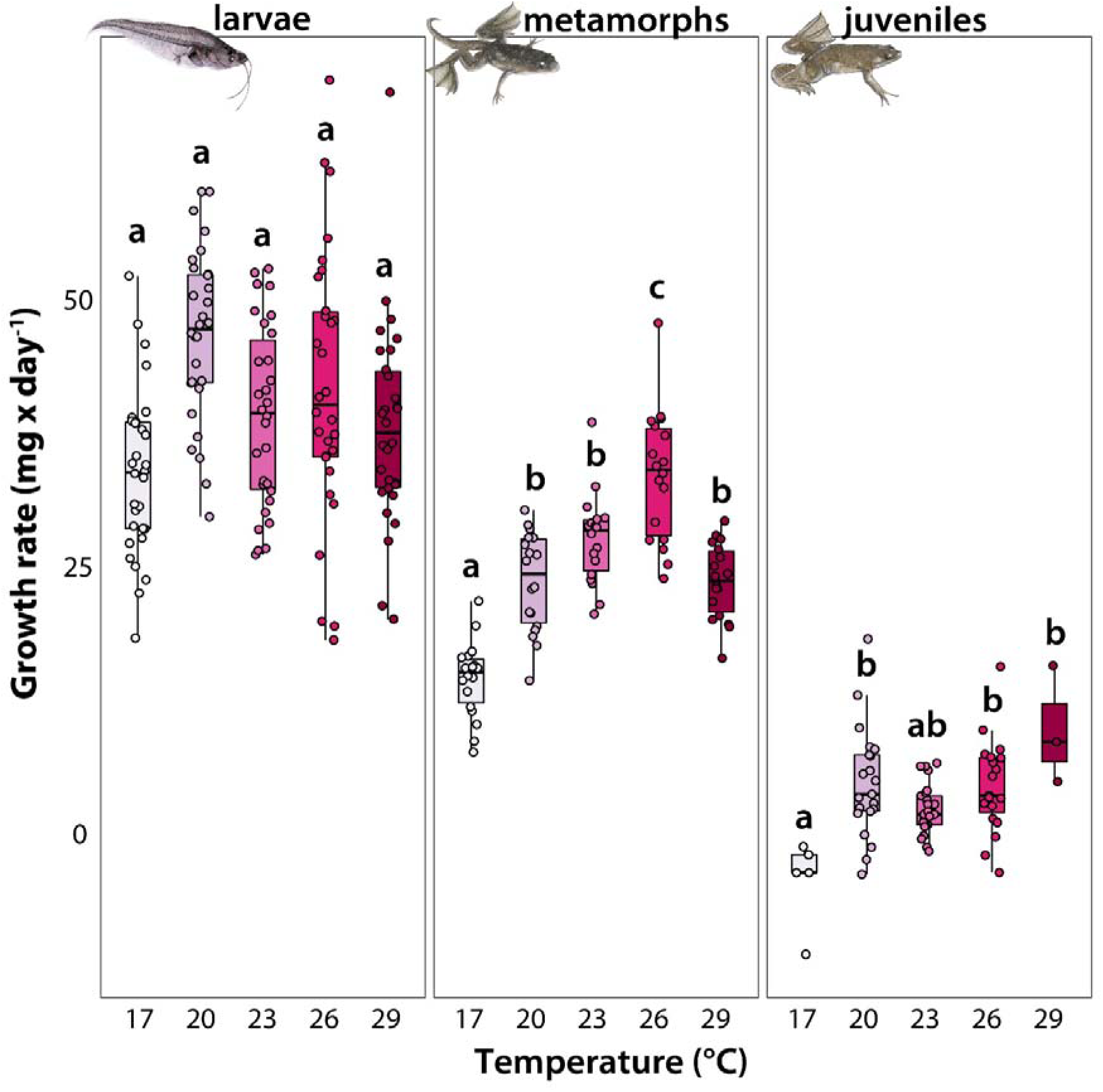
Growth rate (mg x d□¹) of *Xenopus laevis* across three developmental stages (f. l. t. r.: larval, metamorphic, and juvenile) under different rearing temperatures. Each panel represents a distinct life stage, displaying growth rate data for individuals reared at different temperature conditions. Data points represent individual measurements; non-independence among individuals from the same aquarium was accounted for statistically by including tank ID as a random effect in the models.Boxes and whiskers represent the 25^th^ to 75^th^ and 10^th^ to 90^th^ percentiles, respectively; black lines denote medians. Different letters indicate statistically significant differences among temperature treatments within each life stage based on Tukey-adjusted post hoc comparisons (p < 0.05).

*Metamorphs.* Temperature significantly affected SVL, body mass, and growth rate (SVL: χ²=39.56, df=4, p<0.001; mass: χ²=37.39, df=4, p<0.001; growth rate: χ²=77.09, df=4, p<0.001; Table S1). Growth rate increased from 17°C to a peak at 26°C (17 vs. 20, 23, 26, and 29°C all significant, p<0.001–0.007; 20 vs. 26°C, p<0.001; 23 vs. 26°C, p=0.025), before declining significantly at 29°C relative to 26°C (p=0.005). Body mass was significantly lower at 17°C than at 23, 26, and 29°C (p=0.043, p=0.047, p<0.001, respectively), and significantly lower at 20°C than at 29°C (p=0.023); SVL was significantly smaller at 29°C than the other four temperatures (17, 20, 23, and 26°C; all p<0.001–0.039).

*Juveniles.* Temperature significantly affected juvenile growth rate (χ²=6.86, df=1, p=0.0088), although, as for larval growth rate, no individual pairwise comparison remained significant after correction. Temperature did not significantly affect SVL (χ²=0.85, df=1, p=0.358) or body mass (χ²=2.73, df=1, p=0.098).

### 3.2. Heat tolerance and acclimation capacity

CT_max_ was significantly affected by developmental stage (χ²=160.92, df=2, p<0.001), temperature (χ²=153.64, df=4, p<0.001), and their interaction (χ²=60.96, df=8, p<0.001; Table S1), indicating that the shape of the temperature–CT_max_ relationship differed across life stages. Within larvae, CT_max_ increased significantly with temperature across most pairwise comparisons (e.g., 17 vs. 23, 26, and 29°C, all p<0.001). Across stages, juveniles showed significantly higher CT_max_ than larvae at every matched temperature (all p<0.001), and metamorphs showed significantly higher CT_max_ than larvae at most matched temperatures (all p<0.001), consistent with juveniles having the highest heat tolerance overall, followed by metamorphs, then larvae. Body mass did not significantly predict CT_max_ once stage and temperature were accounted for (χ²=0.08, df=1, p=0.782). Acclimation capacity, measured as the ARR for heat tolerance, varied among life stages, being greatest in metamorphs (0.423), intermediate in larvae (0.294), and lowest in juveniles (0.151).

**Figure 2.**
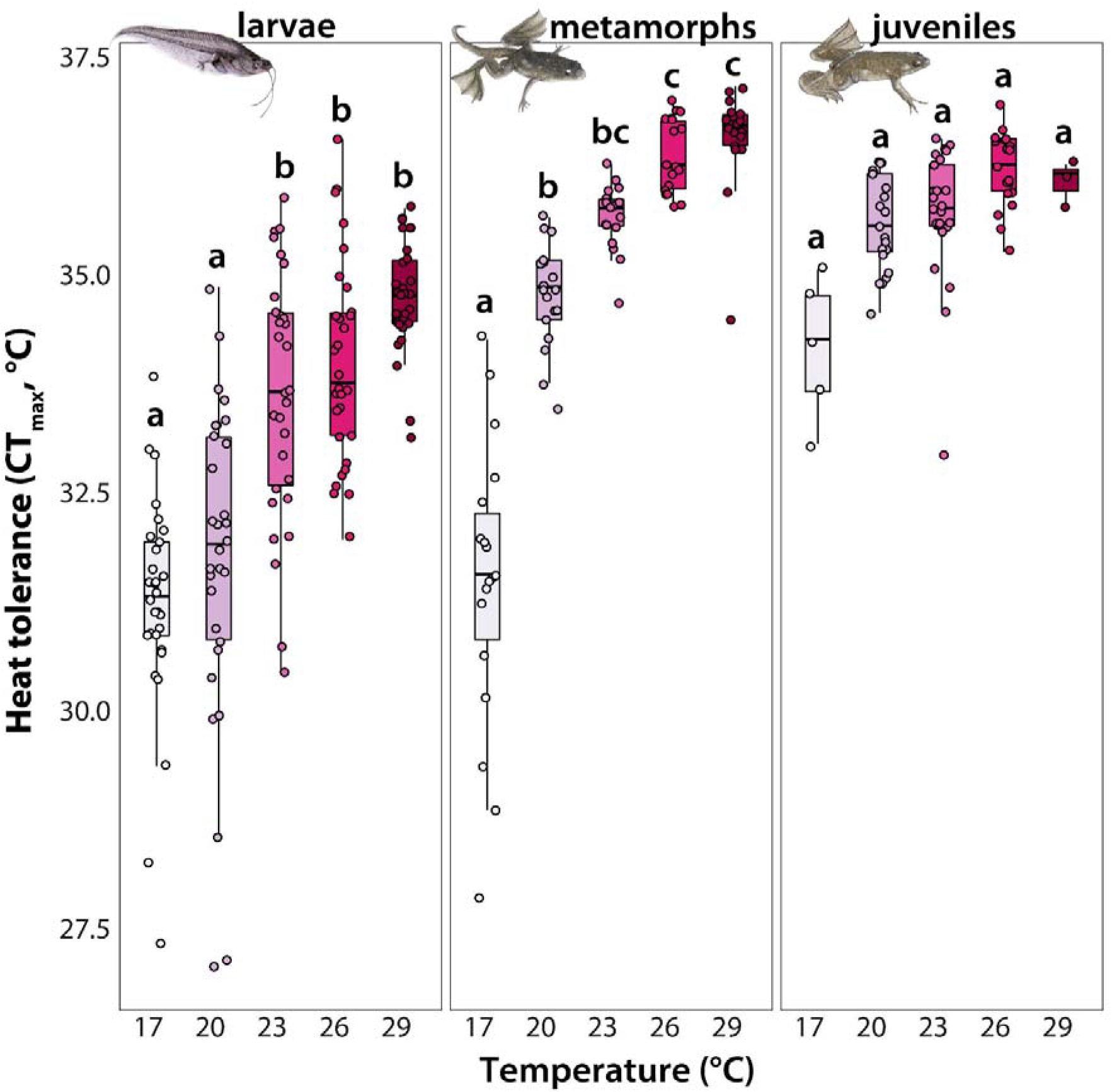
Heat tolerance measured as critical thermal maximum (CT_max_; °C) of *Xenopus laevis* across three developmental stages (f. l. t. r.: larval, metamorphic, and juvenile) under different rearing temperatures. Each panel represents a distinct life stage, displaying heat tolerance data for individuals reared at different temperature conditions. Data points represent individual measurements; non-independence among individuals from the same aquarium was accounted for statistically by including tank ID as a random effect in the models. Boxes and whiskers represent the 25^th^ to 75^th^ and 10^th^ to 90^th^ percentiles, respectively; black lines denote medians. Different letters indicate statistically significant differences among temperature treatments within each life stage based on Tukey-adjusted post hoc comparisons (p < 0.05).

### 3.3. Metabolism

*Routine metabolic rate (RMR)* increased significantly with temperature (χ²=224.24, df=4, p<0.001), with nearly all pairwise comparisons significant (p<0.001–0.007) except between 17°C and 20°C.

*Mean Q_10_*was also significantly affected by temperature (χ²=102.88, df=4, p<0.001), with larvae reared at 26°C and 29°C showing enhanced thermal sensitivity relative to 17–23°C (all p<0.001–0.027), and 29°C differing significantly from 26°C as well (p=0.015).

**Figure 3.**
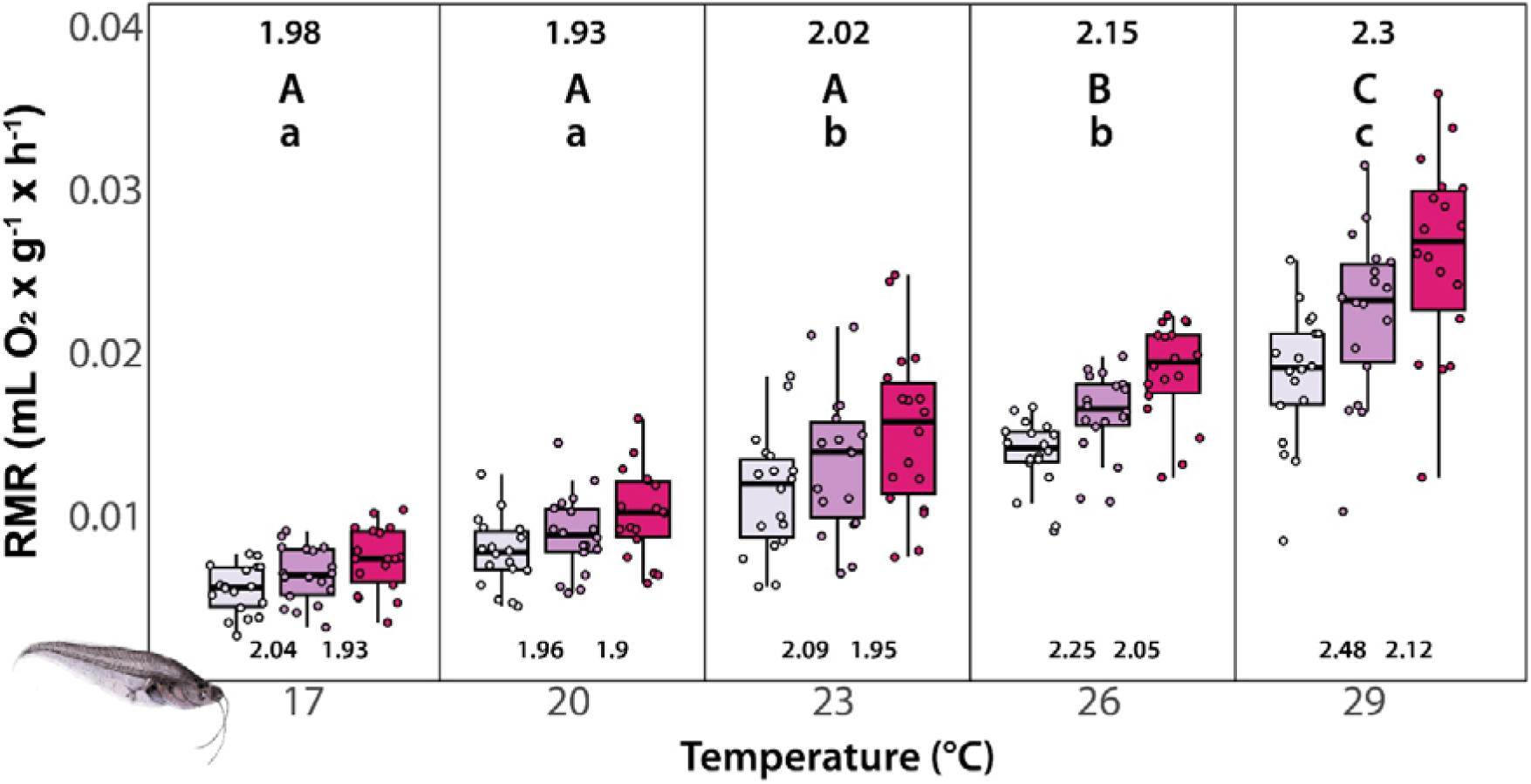
Effect of rearing temperature on routine metabolic rate (RMR, mL O_2_ x g^-1^ x h^-1^) and thermal sensitivity of RMR in larval *Xenopus laevis* measured at developmental stage NF57 (Nieuwkoop et al., 1994). Numbers above the boxes represent the mean thermal sensitivity of RMR, expressed as the average Q_10_ calculated from two sequential steps: step 1, Q between the rearing temperature and rearing temperature +2°C; step 2, Q_10_ between rearing temperature +2°C and +4°C. Numbers below the boxes indicate the individual Q_10_ values for step 1 and step 2, respectively. Boxes and whiskers show 25^th^ to 75^th^ and 10^th^ to 90^th^ percentiles, respectively; black lines indicate the median. Different letters indicate statistically significant differences among temperature treatments within each life stage based on Tukey-adjusted post hoc comparisons (p < 0.05).

### 3.4. CORT release rate

Baseline CORT release rate (i.e., before acute heat stress exposure) was not significantly affected by temperature (χ²=5.70, df=3, p=0.127), with no significant pairwise differences. In contrast, CORT release rate after acute heat stress exposure was significantly affected by temperature (χ²=33.25, df=3, p<0.001), and was significantly higher at 26°C than at both 17°C (p=0.032) and 20°C (p=0.008). Within-treatment (control vs. heatwave) comparisons showed no significant effect at 17°C or 20°C, but significant heatwave-induced increases at 23°C (p=0.013) and 26°C (p=0.0018; Figure 4, S5).

**Figure 4.**
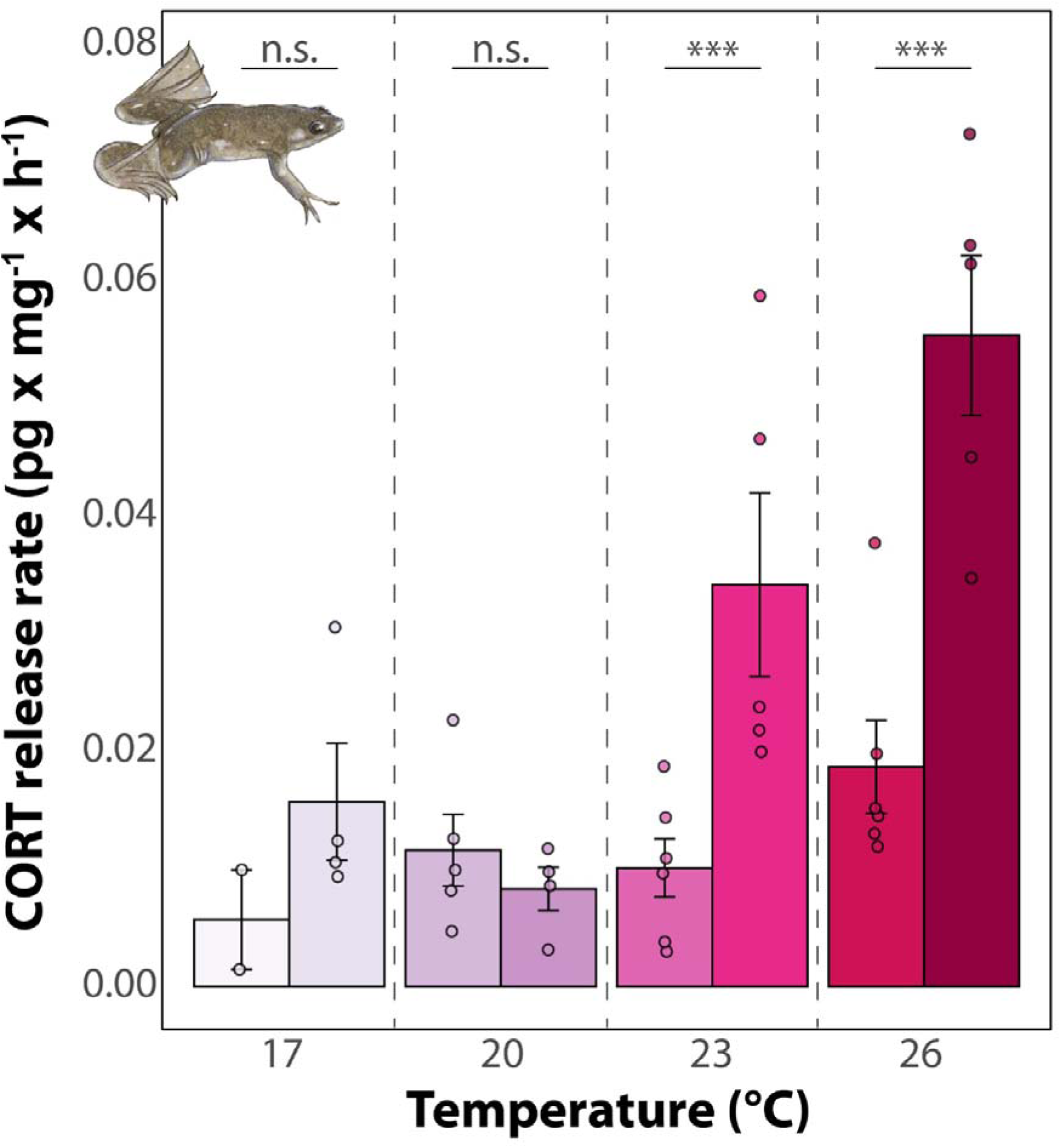
Effect of a 6-day acute heat stress exposure (right bar) on baseline corticosterone (CORT) release rate (pg x mg^-1^ x h^-1^) ±SE in juvenile (150 days after hatching) *Xenopus laevis* compared to specimens maintained at respective rearing temperature (left bar). CORT measures were obtained through waterborne hormone collection. Asterisks indicate statistically significant differences between control and acute heat stress treatments within a temperature group based on within-treatment comparisons (*p < 0.05; **p < 0.01; ***p < 0.001).

### 3.5 Life stage specific fat reserves

Fat reserves were significantly affected by life stage, temperature, and their interaction (Table S1). Metamorphs showed significantly higher relative fat body mass than larvae at every matched temperature (17, 20, 23, and 26°C; all p<0.001–0.029) and higher reserves than juveniles at 17°C (p<0.001). Within metamorphs, fat reserves were significantly higher at 17°C than at 20, 23, and 26°C (all p<0.001), confirming that the highest fat reserves occurred at 17°C. Within larvae, fat reserves were significantly higher at 17°C than at 23°C (p=0.034) and 26°C (p=0.002), but 17°C and 20°C did not differ significantly. No significant pairwise differences were found among juveniles at any temperature.

### 3.6 Survival

Survival showed a non-linear pattern across rearing temperature, but the strength of this pattern differed across developmental stages. Survival to NF57 (larval stage) was not significantly affected by temperature (χ²=1.67, df=4, p=0.796). By NF66 (metamorph stage), temperature had a significant effect (χ²=12.19, df=4, p=0.016), with survival highest at intermediate temperatures and lowest at 29°C, though no individual pairwise comparison remained significant after Tukey correction. By 150 days after hatching, temperature had a strong, significant effect (χ²=80.53, df=4, p<0.001; Firth penalized logistic regression; Table S3), with survival significantly higher at intermediate temperatures than at both extremes (17 vs. 20, 23, and 26°C, all p<0.001; 20, 23, and 26 vs. 29°C, all p<0.001). Separately, the 32°C treatment resulted in complete mortality at the very start of the experiment, before any individuals reached the first sampling stage.

## 4. Discussion

This study investigated whether thermal environments during early life lead to trade-offs between rapid growth and heat tolerance in *X. laevis*, and whether such effects accumulate to influence juvenile resilience to acute thermal stress and survival. We showed that warm rearing temperatures accelerated larval development, maintained growth, and increased heat tolerance, but at the cost of reduced energy reserves, as indicated by smaller fat bodies. Juveniles from warm treatments exhibited elevated CORT responses to acute heat stress and diminished acclimation capacity, indicating higher energetic demands and reduced metabolic flexibility, which may contribute to temperature-driven mortality observed across life stages.

### 4.1 Early-life thermal plasticity promotes both growth and heat tolerance at elevated temperatures

Our study reveals that *X. laevis* larvae reared at elevated temperatures can simultaneously accelerate development, maintain growth, and increase heat tolerance. This pattern contrasts with the prevailing expectation, derived from theories of energy allocation and metabolic trade-offs (Sokolova, 2013, 2018, 2023; Chung & Schulte, 2020), that rapid growth and elevated heat tolerance should compete for limited energetic resources, such that accelerated early-life growth in response to changing environmental conditions would ultimately reduce heat tolerance later in life. Our findings therefore challenge the notion of a strict trade-off between developmental rate and thermal resilience, instead suggesting that amphibians may be able to combine these strategies during early development. Thereby, our results build on previous studies and syntheses showing that developmental thermal environments can shape thermal tolerance and later-life thermal physiology (e.g., Pottier et al., 2022; Ohmer et al., 2023), by demonstrating that these thermal responses occur alongside shifts in life-history traits and energy reserves across larval, metamorphic, and juvenile stages.

Notably, this dual plasticity is likely adaptive in ephemeral habitats. In desiccating ponds, amphibian larvae must balance two opposing risks: the threat of pond drying and the challenge of high diurnal thermal peaks (Griffiths, 1997; Burraco et al., 2023). By accelerating development while simultaneously tolerating heat stress (Drakulic et al., 2016; Enriquez-Urzelai et al., 2019), larvae increase their chances of completing metamorphosis before desiccation (Rudolf & Rödel, 2007), a life-history bottleneck faced by many amphibian species. Such capacity to combine fast development with thermal resilience represents a powerful strategy for coping with the spatiotemporal unpredictability of natural environments, and might be of key importance in an increasingly heterogeneous environment under global change. However, this capacity may depend strongly on resource availability. In our experiment, larvae were fed ad libitum, allowing us to isolate thermal effects without imposing nutritional limitation. In natural habitats, food quantity and quality are likely to vary and may also shift with warming, potentially constraining energy availability for plastic responses and thereby modifying growth, development, and thermal tolerance.

Nevertheless, early-life plasticity often carries potential costs (DeWitt et al., 1998). Accelerated development often results in smaller body size and younger age at metamorphosis (Sinai et al., 2022), both of which are key predictors of future survival, dispersal, and reproductive success in amphibians (rev. in Ruthsatz et al., 2018b). Our results suggest that warm-reared juveniles may compensate for early disadvantages through elevated post-metamorphic growth under persistent thermal conditions, a strategy also observed in other species (Brannelly et al., 2025). However, juvenile size should be interpreted with caution because individuals were sampled at a standardized age after hatching rather than after metamorphosis. Since developmental temperature affected metamorphic timing, treatments differed in the duration of post-metamorphic growth before final sampling. Although juvenile growth rate was calculated relative to time since metamorphosis, juvenile body size at 150 days after hatching therefore integrates developmental temperature effects, metamorphic timing, and post-metamorphic growth duration. While such compensatory growth can partially restore size-related fitness components, it may carry its own costs, including altered morphology, increased oxidative stress, or reduced longevity (Monaghan et al., 2009; Burraco et al., 2020). In our study, these delayed costs were reflected in reduced survival of warm-reared individuals across life stages, particularly at juvenile stage, suggesting that the capacity to exhibit both rapid development and elevated heat tolerance may come at the expense of long-term viability. Thus, the benefits of early-life thermal plasticity in *X. laevis* likely entail subtle but consequential trade-offs that manifest later in the life cycle. In this sense, early developmental conditions may not only shape immediate survival prospects but could also structure life-history trajectories across ontogeny. Understanding how these compensatory dynamics play out under natural thermal regimes will be crucial for assessing the long-term fitness consequences of developmental plasticity in a warming world.

### 4.2 Cumulative effects of elevated temperature during early life constrain resilience to acute heat stress

Exhibiting phenotypic plasticity is energetically costly and often entails trade-offs between maintaining performance under novel environmental conditions and preserving the physiological capacity to respond to additional stressors (DeWitt et al., 1998; Pechenik 1998, 2006). In our study, individuals reared at warmer developmental temperatures consistently exhibited elevated heat tolerance across life stages, indicating that heat tolerance was maintained or even enhanced despite accelerated development. However, our results revealed that this plasticity carries costs. Juveniles from warmer rearing treatments mounted stronger responses of the stress hormone corticosterone (CORT) to acute thermal stress, revealing heightened physiological sensitivity to short-term heat extremes. Notably, baseline CORT levels did not differ among treatments, suggesting that warm-reared individuals were not chronically stressed, but rather had a diminished capacity to modulate their stress response when faced with acute thermal challenges (Alfonso et al., 2020). This pattern implies a trade-off: investment in thermal resilience during development constrains the ability to buffer additional stressors later in life as the higher CORT response might lead to increased energy demands and subsequent cellular damage (Burraco et al., 2020). Supporting this interpretation, warm-reared juveniles exhibited reduced fat body reserves and the lowest acclimation capacity, indicative of increased energetic expenditure and reduced metabolic flexibility (Ruthsatz et al., 2020b).

Because thermal treatments were maintained from larval through juvenile stages, these juvenile responses should not be interpreted as larval-only carry-over effects. Rather, they likely reflect cumulative thermal-history effects, in which larval thermal experience, metamorphic timing, and continued juvenile acclimation jointly shaped later-life stress physiology and survival. Similar costs were apparent in larvae. Individuals reared at higher temperatures showed reduced metabolic stability at elevated test temperatures, expressed as less stable RMRs and higher Q_10_ values. Despite maintaining high heat tolerance, these larvae also showed reduced fat body reserves, indicating that the energetic costs of thermal acclimation are concealed within metabolic regulation rather than reflected in growth metrics. Together, these larval and juvenile responses suggest that sustained thermal exposure across ontogeny can progressively reduce metabolic flexibility and constrain the capacity to sustain energy homeostasis under acute thermal stress.

This interpretation should be considered in light of our controlled thermal design. Constant rearing temperatures allowed us to isolate the effects of sustained thermal exposure, but natural aquatic habitats often experience diel and seasonal thermal fluctuations, including cooler periods that may allow partial physiological recovery. Thus, fluctuating regimes may modify the magnitude of the cumulative costs observed here. Nevertheless, cumulative thermal history remains ecologically relevant, particularly in persistently warm or warming aquatic environments where elevated temperatures may affect individuals across multiple developmental stages. Reciprocal-transfer or common-garden designs would be needed to disentangle larval-only carry-over effects from continued acclimation experimentally.Notably, growing evidence suggests that early-life stress exposure can enhance later resilience to stressful environments. In the context of global change, such hardening, defined as repeated exposure to the same stressor, or cross-tolerance, where exposure to one stressor increases tolerance to another,may act as pre-adaptations (Rodgers & Gomez Isaza, 2022), enabling organisms to better withstand consecutive challenges (Todgham & Stillman, 2013). For instance, Scott & Johnston (2012) found that warming during the embryonic stage improved thermal hardiness later in life in zebrafish (*Danio rerio*). For amphibians this is particularly relevant, as larvae often face thermal or desiccation stress in ponds, while juveniles and adults are exposed to extreme temperature fluctuations in terrestrial habitats. Hardening or cross-tolerance could therefore provide critical resilience across life stages and should be highly adaptive in the face of global change (Rodgers & Gomez Isaza, 2023).

On the contrary, softening or cross-susceptibility can be maladaptive, reducing survival and ultimately threatening fitness and population persistence under global change (Lundsgaard et al., 2023). In our study, development under thermal stress resulted in softening towards acute heat exposure later in life, likely driven by energy depletion or oxidative stress (Burraco et al., 2020; Sokolova, 2013). Similar patterns have been reported in the field cricket *Gryllus lineaticeps* (Harter & Stahlschmidt, 2025). These findings suggest that climate change may pose greater risks to some species than previously recognized. Understanding whether early-life stress exposure enhances or undermines resilience to future challenges is therefore crucial for predicting responses to multi-stressor environments, and identifying the mechanisms underlying these processes remains a key research priority (Rodgers & Gomez Isaza, 2023).

### 4.3 Concluding remarks: Ontogenetic trade-offs might challenge predictions of species resilience under climate change

Our findings suggest that developmental plasticity facilitates coping with elevated mean temperatures but involves trade-offs that compromise physiological buffering capacity under acute thermal extremes. Importantly, larvae reared at warm temperatures were able to accelerate development, maintain growth, and enhance heat tolerance simultaneously, reflecting effective early-life thermal acclimation. Yet, both larvae and juveniles from these treatments were less resilient to acute heat stress, revealing hidden vulnerabilities not apparent from growth or heat tolerance alone. These results underscore that the benefits of developmental plasticity may accumulate as physiological costs across later life stages (Pechenik, 2006), with important implications for predicting species’ resilience under climate change (Lundsgaard et al., 2023).The population studied here had been maintained under constant thermal laboratory conditions for generations. In other taxa, such as zebrafish, long-term exposure to stable environments has been shown to erode phenotypic plasticity because maintaining the genetic and physiological capacity for plastic responses is energetically costly (Morgan et al., 2019, 2022). In contrast, our results establish that *X. laevis,* a fully aquatic and widely invasive species, has demonstrated substantial thermal plasticity across life stages despite its long history under laboratory conditions. This capacity likely contributes to the persistence and spread of invasive populations originating from laboratory escapes or releases, underpinning the species’ success in establishing across diverse habitats and climates. However, while *X. laevis* exhibits substantial thermal plasticity at elevated temperatures, it remains unclear whether this capacity has diminished during prolonged captivity, highlighting the need to investigate wild populations.Our findings reinforce that models of amphibian vulnerability under climate change must move beyond single-stage or single-trait assessment. Focusing solely on traits such as CT may yield overly optimistic estimates of climate resilience, whereas a multi-trait, cross-stage perspective is needed to capture how developmental trajectories, acute stress resilience, and (cumulative) thermal-history effects across life stage effects interact to shape demographic bottlenecks in increasingly extreme climates (Pottier et al., 2025). Likewise, efforts to define thermal thresholds or identify physiological biomarkers of population vulnerability must account for how these traits shift across ontogeny (Ruthsatz & Kruger, 2025). Moving forward, longitudinal, reciprocal-transfer, common-garden, and lifetime-monitoring approaches will be critical for revealing how plasticity unfolds over time and for improving predictions of species persistence under global warming.

## Acknowledgements

We are grateful to Jutta Dammann from the Universitätsklinikum Hamburg Eppendorf for her great commitment in the husbandry and breeding of *X. laevis*. We thank Miguel Vences for his support and Gabriele Keunecke for her help in the laboratory in Braunschweig.

## Author contributions

**IRMM:** Methodology, data curation, formal analysis, investigation (lead), writing – review & editing.

**PB:** Writing – review & editing.

**CH, LK, CM:** Investigation, data curation, writing – review & editing.

**NK:** Conceptualization, methodology, formal analysis (lead), writing – original draft, writing – review & editing.

**KR:** Conceptualization, supervision, methodology, data curation, formal analysis, investigation, writing – original draft (lead), writing – review & editing.

## Conflict of Interest

The authors declare that the research was conducted in the absence of any commercial or financial relationships that could be construed as a potential conflict of interest

## Data availability

Data, including all original measurements, and codes will be deposited in Figshare and Github after acceptance.

## Statement of Ethics

Breeding was conducted under permission from the *Behörde für Justiz und Verbraucherschutz Hamburg*, Germany (Gz. N111/2023). The experiments were conducted under permission from the *Niedersächsisches Landesamt für Verbraucherschutz und Lebensmittelsicherheit*, Germany (Gz. 33.19-42502-04-23-00492).

## Funding

This study was funded by intramural funds of the Technische Universität Braunschweig. IRMM was supported by the Company of Biologist Travelling Fellowship (No. JEBTF23101266), the Fundação Coordenação de Aperfeiçoamento de Pessoal de Nível Superior (CAPES, No. 88887.700216/2022-0) scholarship, and Fundação Carlos Chagas Filho de Amparo à Pesquisa do Estado do Rio de Janeiro (FAPERJ), Grant Nota 10, Process N°. E-26/201.637/2025. KR was supported by a Marie Curie Fellowship AMPHISTRESS-101151070.The German Research Foundation (DFG) project (No. 459850971; *A new actor on the stage of global change: A multi-level perspective on the toxicity of microplastics pollution in amphibians*) supported CM and KR. PB was supported by Ramón y Cajal fellowship 2023-044964-I (Spanish Ministry of Science and Innovation). NK was supported by Dr. Marie Wasdell & Malkito Kaur Gangar Fellowship (*Investigating the ecophysiology of the globally invasive African clawed frog, Xenopus laevis*).

## Notes

### Competing Interest Statement

The authors have declared no competing interest.

### Summary of Updates

We have carefully revised the manuscript and addressed all points raised. The main changes include a more detailed presentation of the statistical analyses, revised figures with significance annotations, clarification of several methodological aspects, and a more explicit discussion of the limitations and ecological interpretation of our experimental design. We have also substantially revised the statistical approach and provide full model outputs and pairwise comparisons in the Supporting Information.

