## Supplementary Material for "Ontogenetic consequences of developmental temperature in amphibians: simultaneous gains in heat tolerance and cumulative costs to stress physiology"

**Supporting Information**

**Glossary**

**Thermal tolerance:** The range of environmental temperatures within which an organism can sustain essential physiological functions, bounded by lower and upper critical limits. This range is often illustrated by thermal performance curves, which describe how physiological traits such as metabolism, growth, or locomotion vary with temperature, peaking at an optimal temperature (T_Opt_) and declining toward the organism’s critical thermal limits (CT_min_ and CT_max_).

**Critical thermal limit (CT):** The extreme temperatures (both minimum, CT_min_, and maximum, CT_max_) beyond which an organism loses essential physiological functions, such as locomotion or metabolic stability, leading to potential mortality.

**Heat tolerance:** An organism’s ability to withstand elevated temperatures, often quantified by the critical thermal maximum CT_max_, which is the highest temperature at which survival is possible.

**Optimal temperature (T_Opt_):** The specific temperature at which an organism's physiological processes, such as enzyme activity or metabolic rate, operate at their highest efficiency, defined as the highest functional output per unit time (e.g., maximal net energy gain for metabolism or maximal catalytic turnover for enzymes) before performance declines due to reduced stability or thermal stress.

**Thermal stress**: Thermal stress occurs when temperatures deviate substantially from an organism’s optimum temperature (T_Opt_), causing reduced metabolic efficiency, impaired enzymatic function, disruption of cellular homeostasis, and increased energetic demands. Prolonged or intense thermal stress can limit growth, compromise immune and endocrine function, reduce survival, and ultimately push individuals toward their critical thermal limits (CT_min_ or CT_max_).

**Thermal sensitivity:** The degree to which an organism's physiological traits or performance metrics change in response to variations in environmental temperature.

**Acclimation capacity:** The ability of an organism to physiologically adjust to changes in environmental temperature over time through reversible modifications. This capacity represents a form of phenotypic plasticity, which may manifest as changes in thermal tolerance, metabolic rate, or other physiological traits.

**Thermal reaction norm:** A graphical representation depicting how a specific trait or performance measure of an organism varies across a range of temperatures, illustrating its phenotypic plasticity in response to thermal environments.

**Heat wave:** A heat wave is a prolonged period of unusually high temperatures, often accompanied by high humidity, that significantly exceeds the average for a given location and time of year. These events are often accompanied by elevated humidity, which can exacerbate thermal stress by impairing evaporative cooling and increasing the physiological heat load experienced by organisms.

**Material and Methods**

*Study species*

The African clawed frog, *Xenopus laevis*, was selected as a model organism because it is ecologically important as a globally invasive amphibian (Lillo et al., 2011; Measey et al., 2012; Courant et al., 2018; Mora et al., 2019; Peñafiel-Ricaurte et al., 2023) and exhibits resilience to environmental change (Wilson et al., 2000, Araspin et al., 2020, Wagener et al., 2021, Kruger et al., 2022). Additionally, it experiences a wide range of temperatures and tadpoles display a broad 80% thermal performance breadth (13.1°C to 36.7°C) in their native range in South Africa (Ginal et al., 2023), allowing us to experimentally test how early-life thermal environments influence subsequent thermal resilience and development. Furthermore, this species is a well-established laboratory model due to its trade for use in developmental studies (Carotenuto et al., 2023). We used a captive population of *X. laevis* maintained under standard laboratory conditions. The stable and moderate conditions of laboratory facilities may affect the acclimation ability and heat tolerance as plasticity may erode in these conditions as it is costly to maintain (De Witt et al., 1998; Crates et al., 2023). While captive populations may differ from wild conspecifics in certain ecological and physiological traits (Turko et al., 2023), understanding the thermal plasticity of lab-maintained lineages is relevant for assessing growth-thermal tolerance trade-offs as they are a major source of invasive *Xenopus laevis* populations globally.

*Life history variables*

We measured snout–vent length (SVL) and body mass at stage NF45 (start of experiment, 6 dah) in 20 randomly selected larvae per clutch (N = 60) to calculate growth rate, and again at the three sampling points (i.e., NF57= larvae; NF66= metamorphs; 150 dah = juveniles) (Fig. S1). Snout-vent-length was measured to the nearest 0.5 mm from photographs taken on laminated graph paper after terminal sampling, using ImageJ (Schneider et al., 2012). To measure body mass, specimens were dry blotted and weighed to the nearest 0.001 g with an electronic balance (Sartorius A200 S, Germany). Larval and metamorph growth rates (mg × d^-1^) were calculated from mass at respective sampling points minus the mass at stage NF45, divided by the days from stage 45 to the sampling point. Juvenile growth rate (mg × d^-1^) was calculated from individual mass at the respective sampling point (i.e., 150 dah) within a temperature treatment minus the mean mass of metamorphs within a temperature treatment, divided by the age in days since the start of the experiment minus the mean age of juveniles at completion of metamorphosis in days.

We documented the date of the first and last animal to complete metamorphosis per aquarium to account for within-treatment variation in developmental rate. At 150 days after hatching, animals from different temperature treatments had completed metamorphosis 59.3±9.71 (17°C), 100.83±1.65 (20°C), 110.22±1.76 (23°C), 117±1.18 (26°C), and 104.22±3.74 (29°C) ago.

*Heat tolerance assessments*

For heat tolerance assessments, each individual was placed in a 250 mL beaker with 150 mL of aerated water and submerged in a temperature-controlled water bath set to the individual’s acclimation temperature. Water temperature was increased at +0.1 °C min⁻¹ (Lutterschmidt & Hutchison 1997). Upon reaching CT_max_, animals were transferred to a 1 L beaker at a recovery temperature calculated as

$T=acclimation temperature+ \frac{(CTmax- acclimation temperature)}{2}$,

and placed in a water bath at the rearing temperature for gradual cooling. Once fully recovered (normal swimming), individuals were anaesthetized in buffered tricaine methanesulfonate (MS-222; larvae: 2 g L⁻¹; metamorphs and juveniles: 6 g L⁻¹) until unresponsive to stimuli. Body mass (dry blotted) and SVL were measured, the left side (larvae) or dorsal side (metamorphs, juveniles) was photographed, and specimens were snap-frozen in liquid nitrogen and stored at −80 °C for later fat body dissections and DNA extractions. Trials were conducted between 10:00 and 20:00 h to control for diel variation and match natural peak temperatures (Agudelo-Cantero & Navas, 2019).

*Metabolism measurements*

To measure oxygen consumption, larvae were placed individually in 30 mL glass vials filled with autoclaved tap water to prevent microbial oxygen consumption and sealed with airtight rubber stoppers. A chemical-optical oxygen sensor spot was integrated into each vial and connected to a multi-channel oxygen measurement system (Oxy-4 SMA; PreSens Precision Sensing GmbH, Regensburg, Germany) via a fiber optic sensor (Polymer Optical Fiber POF, PreSens Precision Sensing GmbH, Regensburg, Germany). A temperature probe immersed in water at the same temperature as the respiratory vials provided automatic temperature compensation for dissolved oxygen measurements.

Before each trial, sensors were calibrated using air-saturated water for the 100% reference point and a factory-set zero oxygen calibration point. The water bath containing the vials was continuously mixed to ensure uniform temperature distribution. Measurements started 10 min after introducing the animals into the vials to allow acclimation, and oxygen concentration (ml O₂ L⁻¹) was recorded every 15 s for 30 min. Control vials without animals were run concurrently, and their oxygen consumption values were subtracted from experimental readings. To avoid hypoxia-induced changes in metabolism, oxygen levels in each vial were maintained above 80% of the initial oxygen saturation throughout the trial.

To assess acute thermal sensitivity, oxygen consumption was measured not only at the acclimation temperature but also at +2 °C and +4 °C above acclimation temperature (Sinai et al., 2024). The water bath was heated at a rate of +0.1 °C min⁻¹ to the target temperature. During ramping, vials were left open for aeration. Once the target temperature was reached, animals were given a 10 min acclimation period before the 30 min measurement. This procedure was repeated for the second temperature step. After the final measurements, the water bath temperature was gradually decreased to the acclimation temperature at –0.1 °C min⁻¹.

*Thermal sensitivity of RMR*

Sensitivity of RMR to short-term temperature variation (i.e., acute thermal sensitivity) was identified by calculating the coefficient Q_10_, which is a measure of the rate of change of the metabolic processes by an increase of temperature of 10 °C (e.g., Dalvi et al., 2009). The relative change in respiration rate over a 10 °C interval (Q_10_) was calculated using the formula:

$\boldsymbol{Q}\boldsymbol{10=}{\boldsymbol{(}\frac{\boldsymbol{Rate 2}}{\boldsymbol{Rate 1}}\boldsymbol{)}}^{\boldsymbol{(}\frac{\boldsymbol{10}}{\boldsymbol{Temperature 2-Temperature 1}}\boldsymbol{)}}$**,**

where *Rate₁* and *Rate₂* are respiration rates at temperatures *T₁* and *T₂*, respectively. Higher Q₁₀ values indicate greater proportional increases in RMR with rising temperature.

*CORT release and quantification*

At the end of the heat stress resilience assessments (Fig. S1), each animal was gently transferred to a freshly cleaned (1× ethanol, 3× water rinse) 250 mL glass beaker containing 50 mL of aged, filtered, aerated tap water pre-heated to its treatment-specific temperature. Beakers were placed in a water bath to maintain constant temperature throughout the 1 h sampling period. . After the hour-long sample collection period, specimens were removed, and the water sample was collected. For each sampling batch, a control beaker (no animal) was run to detect any background hormonal traces; control samples were pooled prior to extraction. Immediately after sampling, each animal was anaesthetized in 50 mL of 6 g L⁻¹ buffered tricaine methanesulfonate (MS-222), blotted dry, weighed to the nearest 0.001 g, and snap frozen in a sterile 5 mL tube in liquid nitrogen for later fat body dissection. Of the 46 surviving animals, CORT samples were collected from 43; three were excluded due to prolonged capture attempts that could have artificially elevated CORT release.

Water samples were stored at −20 °C and processed within one week. Thawed samples were first filtered through Q8 Whatman filter paper to remove particles and feces, then extracted via C18 solid-phase extraction columns (Oasis Vac Cartridge HLB 3 cc, 60 mg, 30 µm; Waters Inc.) using a vacuum manifold (Visiprep; Sigma-Aldrich). Cartridges were pre-cleaned with 4 mL HPLC-grade ethanol and 4 mL nanopure water before each use. Hormones were eluted with 4 mL HPLC-grade dimethyl ether, transferred into 5 mL Eppendorf tubes, and evaporated under a fine N₂ stream at 45 °C using a sample concentrator and block heater (Stuart SBHCONC/1; SBH130D/3). Dried samples were reconstituted in 125 µL (5% lab-grade ethanol, 95% EIA buffer).

CORT concentrations were measured using the DetectX Corticosterone ELISA kit (K014-H5; Arbor Assays, Ann Arbor, MI, USA), previously validated for *Xenopus laevis* (Ruthsatz et al., 2023). We used the 50 µL format for both standards and samples, running all in duplicate on 96-well plates. Plates were read at 450 nm with a Tecan Spark® Microplate Reader (Tecan, Switzerland). Concentrations were calculated from 4PLC standard curves using MyAssays online tools. Mean duplicate CV was 6.61%, and the average R² for curve fits was 0.999. Following Gabor et al., (2013), CORT release rates (pg mL⁻¹) were multiplied by the resuspension volume (0.125 mL) and standardized to individual body mass, yielding final rates in pg g⁻¹ h⁻¹.

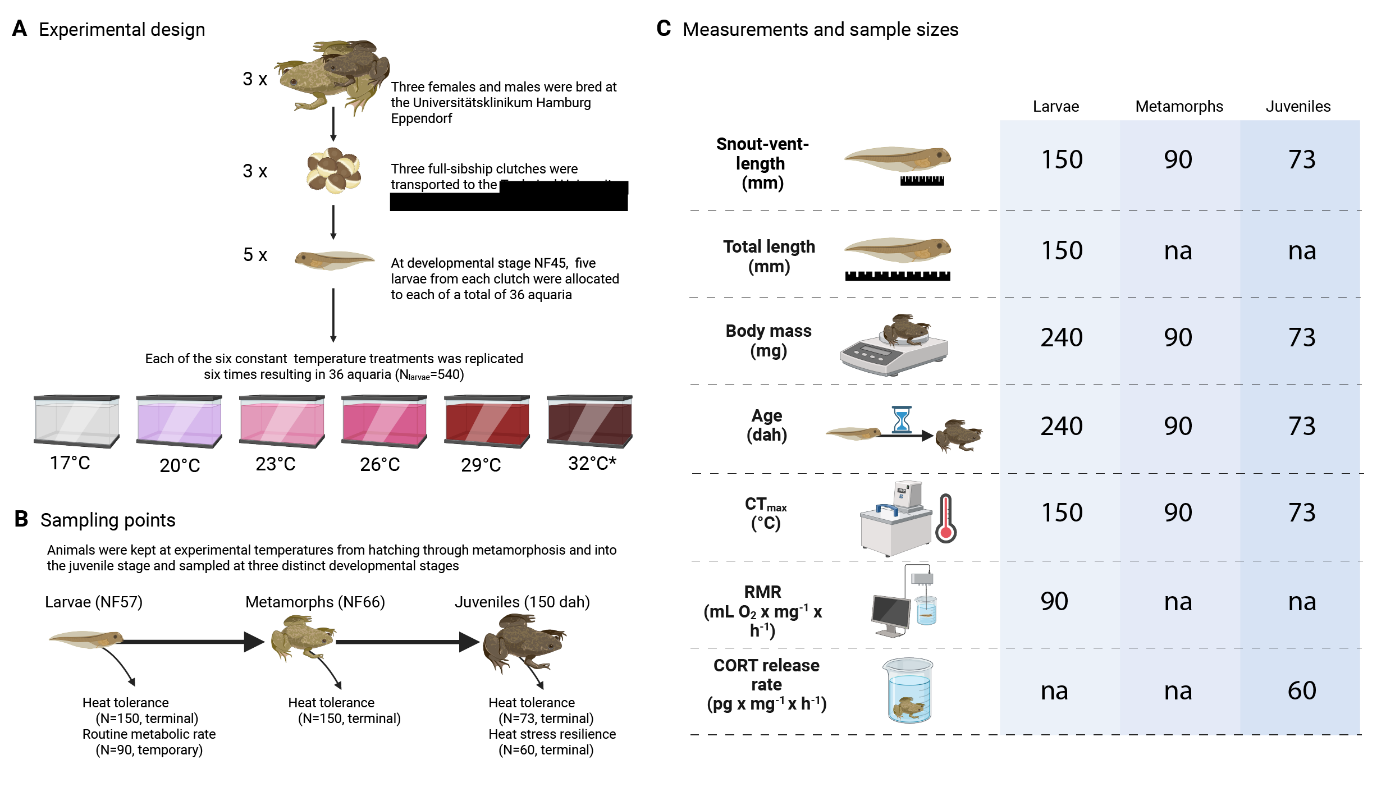
**Figure S1**. Experimental overview with details on **A**. Experimental design, **B**. Sampling points, and **C**. Measurement types and sample sizes. Figure created with Biorender.

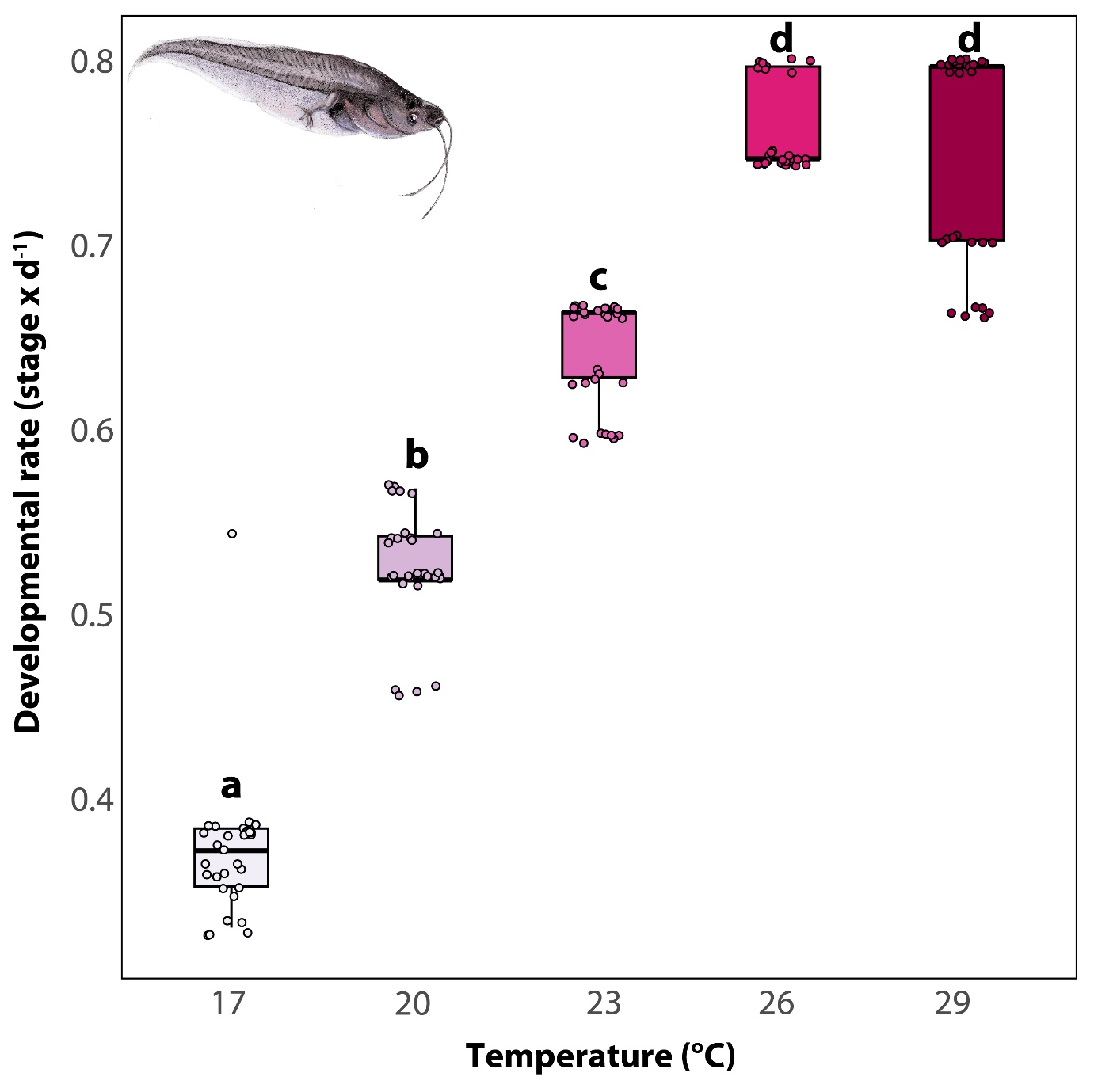
**Figure S2.** Larval developmental rate of *Xenopus laevis* under different rearing temperatures. Data points within each panel correspond to individual measurements, with boxplots summarizing the distribution (median, interquartile range, and range) for each temperature treatment. Boxes and whiskers represent the 25^th^ to 75^th^ and 10^th^ to 90^th^ percentiles, respectively; black lines denote medians.

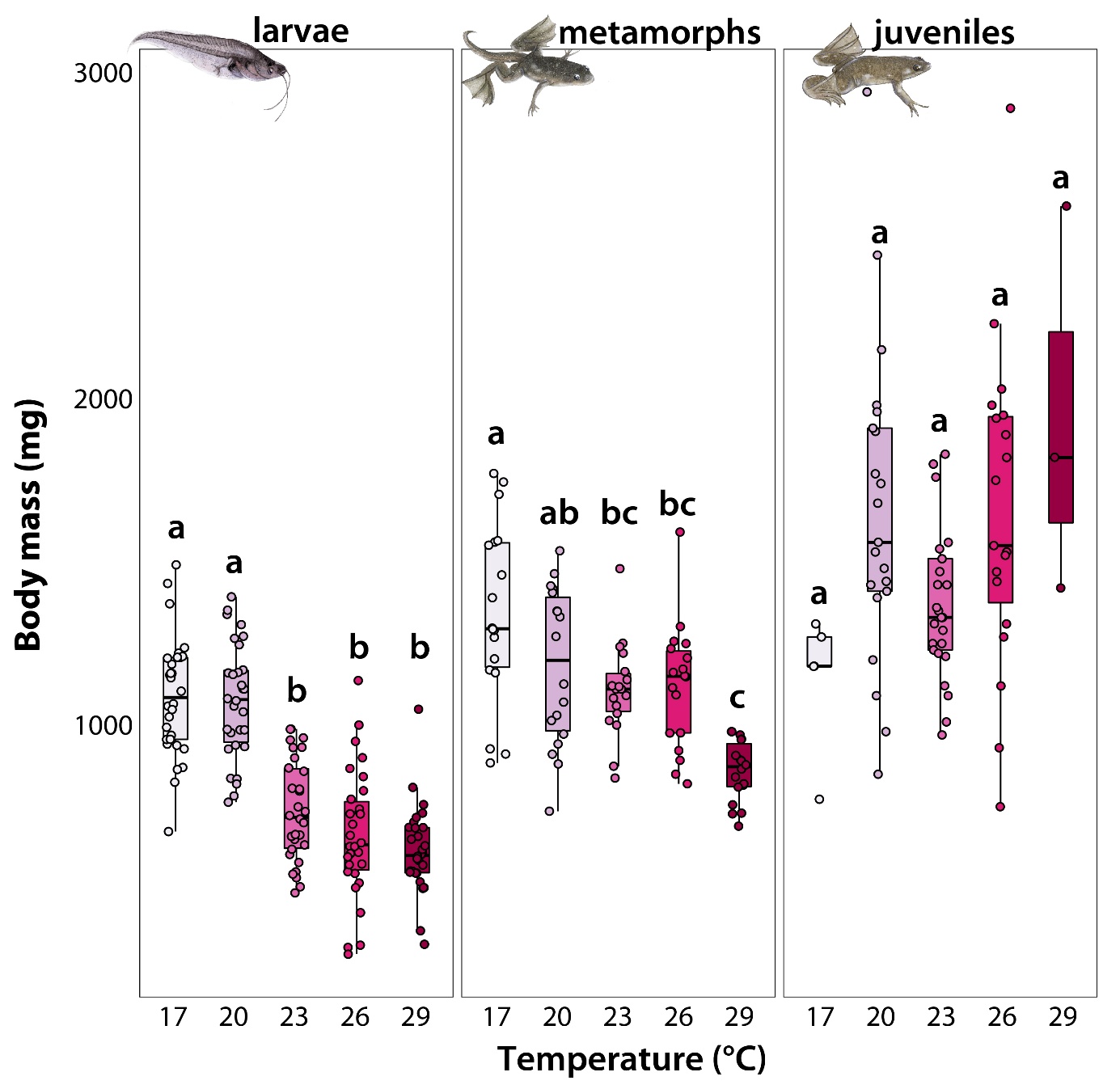
**Figure S3.** Body mass of *Xenopus laevis* across three developmental stages (larval, metamorphic, and juvenile) under different rearing temperatures. Each panel represents a distinct life stage, displaying body mass data for individuals reared at different temperature conditions. Data points within each panel correspond to individual measurements, with boxplots summarizing the distribution (median, interquartile range, and range) for each temperature treatment. Boxes and whiskers represent the 25^th^ to 75^th^ and 10^th^ to 90^th^ percentiles, respectively; black lines denote medians.

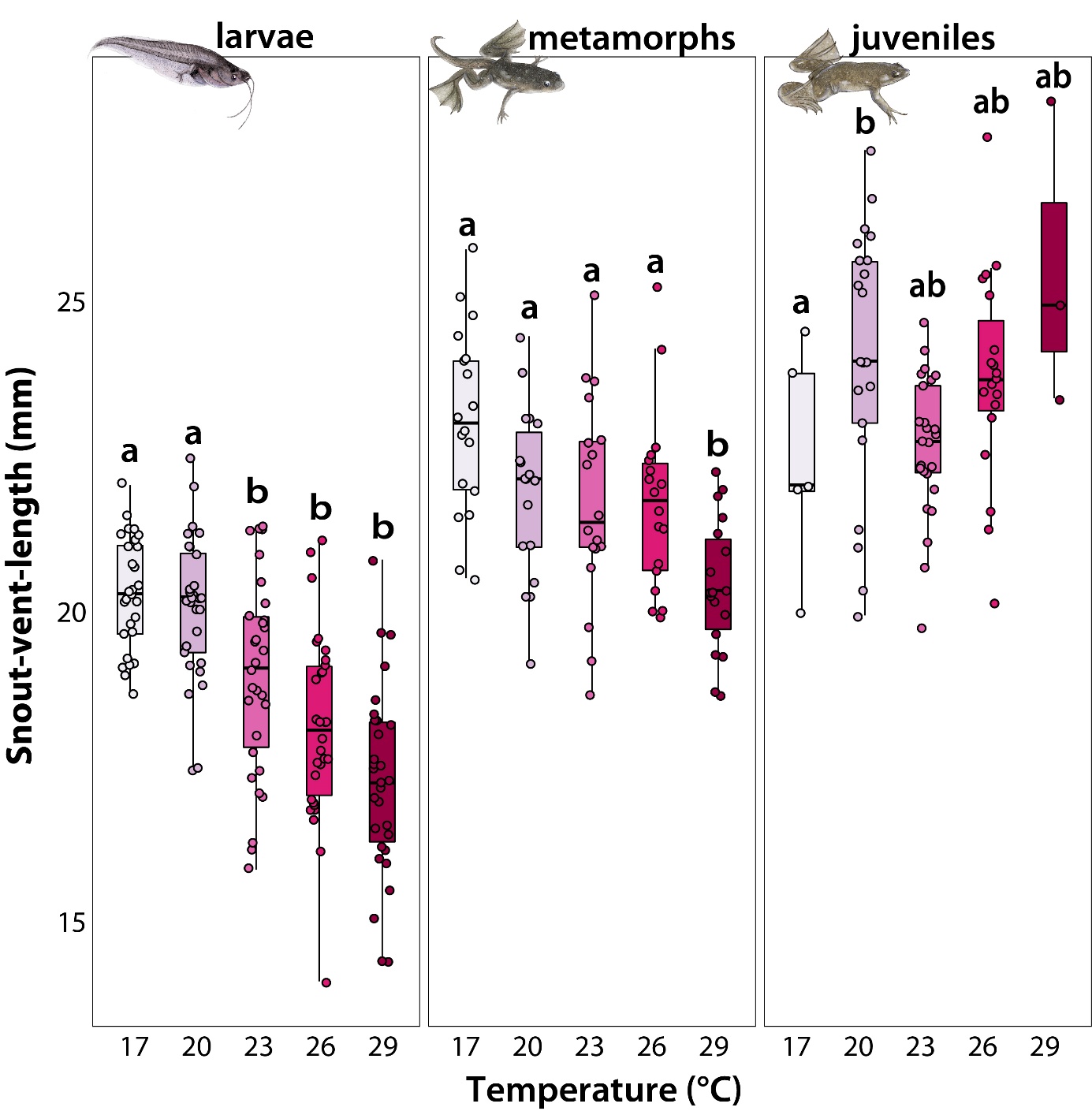
**Figure S4.** Snout-vent-length (SVL) of *Xenopus laevis* across three developmental stages (larval, metamorphic, and juvenile) under different rearing temperatures. Each panel represents a distinct life stage, displaying SVL data for individuals reared at different temperature conditions. Data points within each panel correspond to individual measurements, with boxplots summarizing the distribution (median, interquartile range, and range) for each temperature treatment. Boxes and whiskers represent the 25^th^ to 75^th^ and 10^th^ to 90^th^ percentiles, respectively; black lines denote medians.

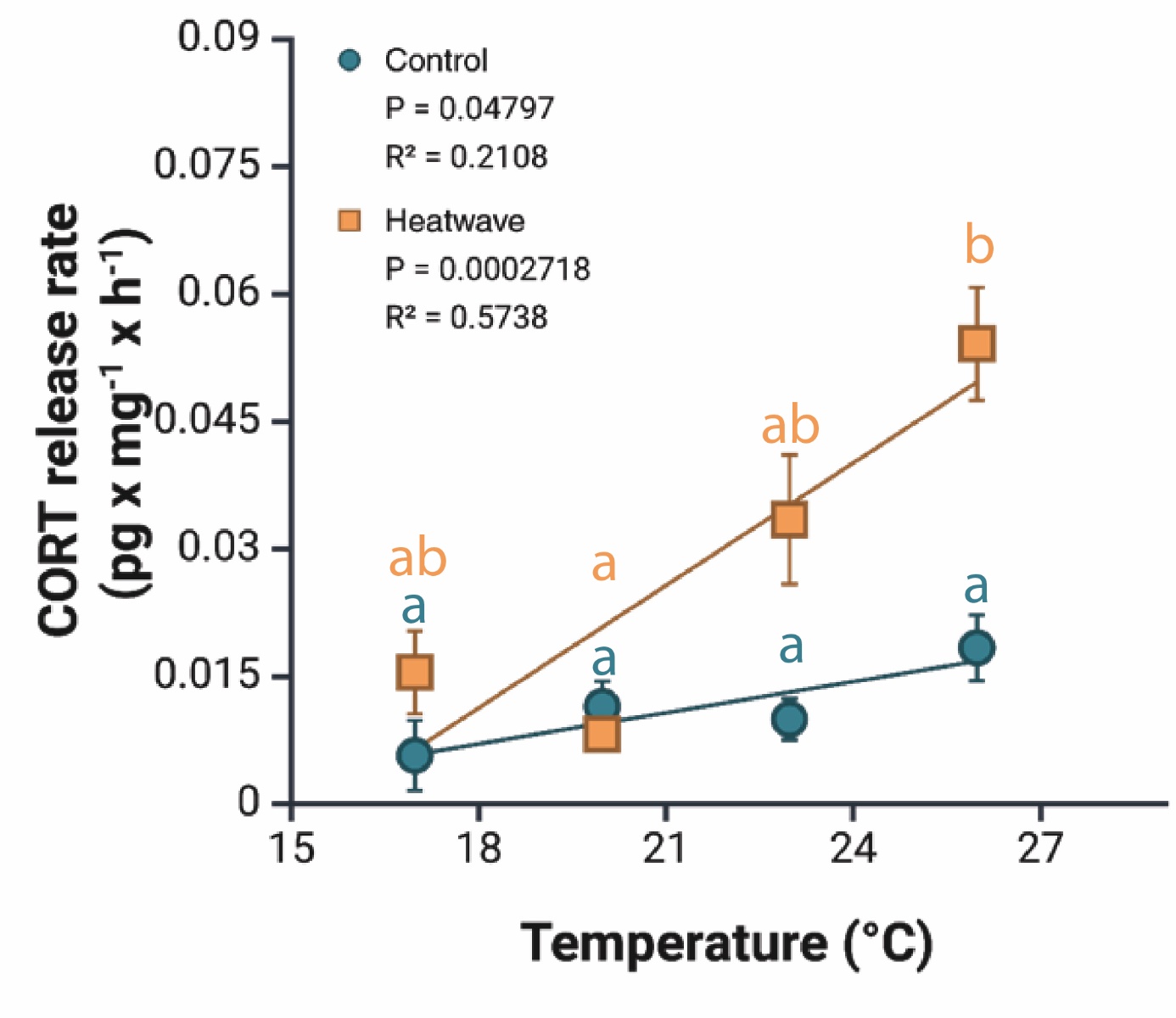
**Figure S5**. Linear regression of rearing temperature and CORT release rate (pg x mg^-1^ x h^-1^) in juvenile *Xenopus laevis* (150 days after hatching) exposed to either control (i.e., rearing temperature; green line and dots) or heatwave (rearing temperature + 5°C; orange line and squares) conditions for 6 days. Shown are mean values ± SE.

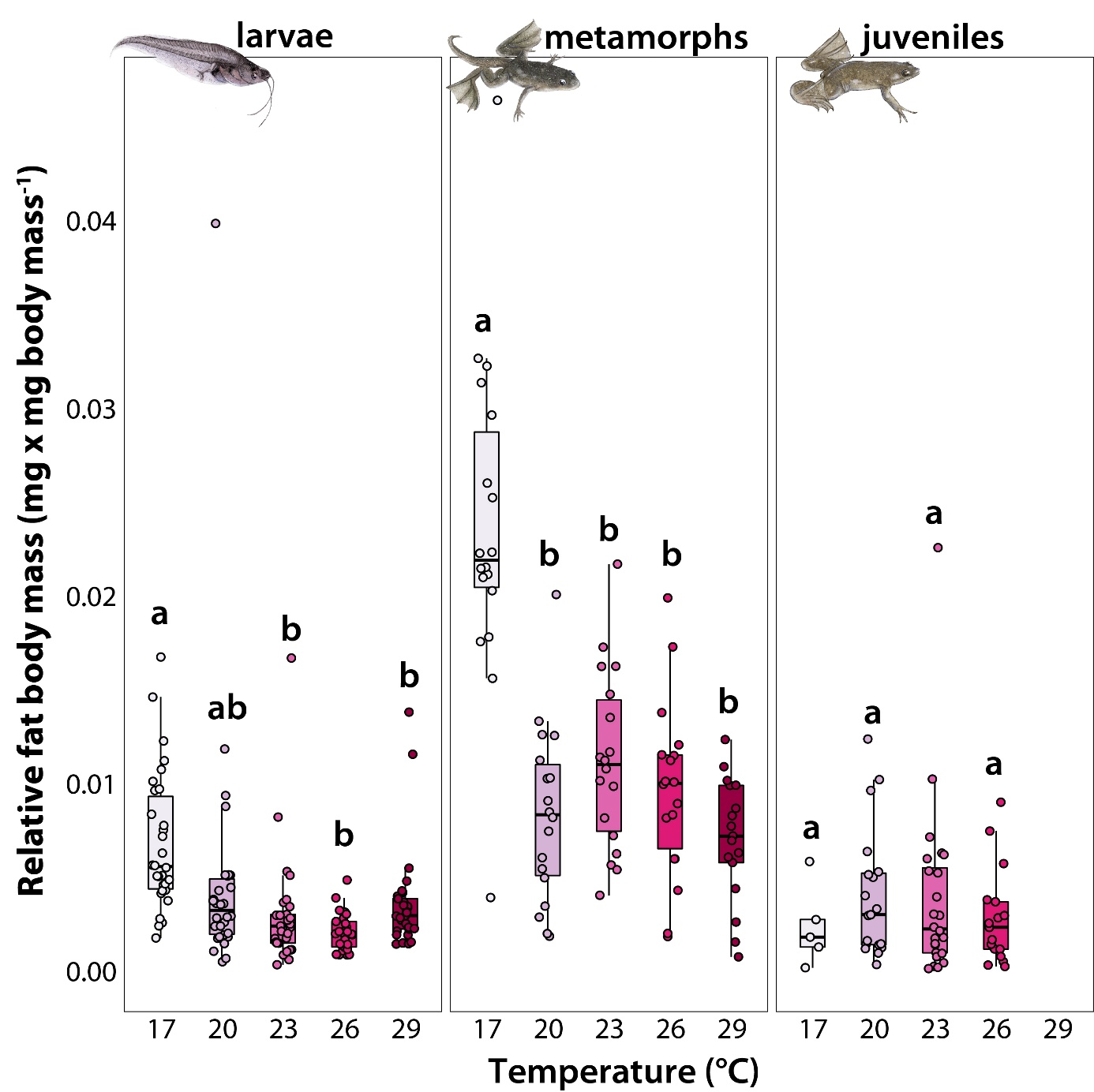
**Figure S6.** Relative fat body size (mg x mg⁻¹ body mass) of *Xenopus laevis* across three developmental stages (larval, metamorphic, and juvenile) under different rearing temperatures. Each panel represents a distinct life stage, displaying relative fat body size data for individuals reared at different temperature conditions. Data points within each panel correspond to individual measurements, with boxplots summarizing the distribution (median, interquartile range, and range) for each temperature treatment. Boxes and whiskers represent the 25^th^ to 75^th^ and 10^th^ to 90^th^ percentiles, respectively; black lines denote medians.

**Table S1:** Results of mixed-effects models testing the effects of rearing temperature and, where applicable, tank-level survival rate (a proxy for density), life stage, or an acute heatwave challenge, on developmental, growth, size, physiological, and survival traits across larval, metamorph, and juvenile life stages of *Xenopus laevis*. Gaussian models included Tank (or Replicate) as a random intercept; the survival model for 150 days after hatching used Firth penalized logistic regression without a random effect, following complete separation in the corresponding mixed model (see Methods). Stage indicates the life stage, Dependent variable is the response variable modelled; Independent variable is the fixed-effect predictor tested, with models containing more than one fixed effect shown across multiple rows, one per predictor. Model family indicates the distribution family and modelling framework used: Gaussian linear mixed models (fitted via lmer), binomial generalized linear mixed models (fitted via glmer), and Firth penalized logistic regression (logistf). Chi-sq is the Wald chi-square test statistic for the given predictor (or, for the Firth model, the penalized likelihood-ratio statistic); Df is the associated degrees of freedom; p-value is the significance of the predictor's effect. Marginal R² is the proportion of variance explained by the fixed effects alone, and Conditional R² additionally includes the random effect, where estimable (NA where the random effect variance could not be reliably computed, including for the Firth model, which has no random effect). Significant effects (p < 0.05) are shown in bold.

| Stage | Dependent variable | Independent variable | Model family | Chi-sq | Df | p-value | Marginal R2 | Conditional R2 |
| --- | --- | --- | --- | --- | --- | --- | --- | --- |
| Larvae | Developmental rate (dah as proxy) | Temperature | Gaussian (lmer) | 669.17 | 4 | **<0.001** | 0.936 | 0.973 |
|  | SVL | Temperature | Gaussian (lmer) | 79.33 | 4 | **<0.001** | 0.484 | 0.573 |
|  |  | Survival rate | Gaussian (lmer) | 2.53 | 1 | 0.111 | 0.484 | 0.573 |
|  | Body Mass | Temperature | Gaussian (lmer) | 251.51 | 4 | **<0.001** | 0.636 | 0.67 |
|  |  | Survival rate | Gaussian (lmer) | 1.76 | 1 | 0.185 | 0.636 | 0.67 |
|  | Growth rate | Temperature | Gaussian (lmer) | 9.81 | 4 | **0.0438** | 0.224 | 0.456 |
|  |  | Survival rate | Gaussian (lmer) | 5.26 | 1 | **0.0218** | 0.224 | 0.456 |
|  | RMR | Temperature | Gaussian (lmer) | 224.24 | 4 | **<0.001** | 0.718 | NA |
|  | Q10 | Temperature | Gaussian (lmer) | 102.88 | 4 | **<0.001** | 0.536 | NA |
| Metamorph | SVL | Temperature | Gaussian (lmer) | 39.56 | 4 | **<0.001** | 0.429 | 0.569 |
|  |  | Survival rate | Gaussian (lmer) | 0.41 | 1 | 0.523 | 0.429 | 0.569 |
|  | Body Mass | Temperature | Gaussian (lmer) | 37.39 | 4 | **<0.001** | 0.379 | 0.495 |
|  |  | Survival rate | Gaussian (lmer) | 0.8 | 1 | 0.37 | 0.379 | 0.495 |
|  | Growth rate | Temperature | Gaussian (lmer) | 77.09 | 4 | **<0.001** | 0.659 | 0.776 |
|  |  | Survival rate | Gaussian (lmer) | 2.26 | 1 | 0.133 | 0.659 | 0.776 |
| Juvenile | SVL | Temperature | Gaussian (lmer) | 0.85 | 1 | 0.358 | 0.022 | 0.278 |
|  | Mass_mg | Temperature | Gaussian (lmer) | 2.73 | 1 | 0.0982 | 0.068 | 0.318 |
|  | Growth rate | Temperature | Gaussian (lmer) | 6.86 | 1 | **0.00884** | 0.177 | 0.491 |
| All stages | CTmax | Stage | Gaussian (lmer) | 160.92 | 2 | **<0.001** | 0.732 | 0.769 |
|  |  | Temperature | Gaussian (lmer) | 153.64 | 4 | **<0.001** | 0.732 | 0.769 |
|  |  | Body mass | Gaussian (lmer) | 0.08 | 1 | 0.782 | 0.732 | 0.769 |
|  |  | Stage*Temperature | Gaussian (lmer) | 60.96 | 8 | **<0.001** | 0.732 | 0.769 |
|  | Life stage specific fat reserves | Stage | Gaussian (lmer) | 199.34 | 2 | **<0.001** | 0.578 | 0.578 |
|  |  | Temperature | Gaussian (lmer) | 43.23 | 3 | **<0.001** | 0.578 | 0.578 |
|  |  | Growth rate | Gaussian (lmer) | 6.62 | 1 | **0.0101** | 0.578 | 0.578 |
|  |  | Stage*Temperature | Gaussian (lmer) | 63.82 | 6 | **<0.001** | 0.578 | 0.578 |
| Baseline (Control) | CORT release rate | Temperature | Gaussian (lmer) | 5.7 | 3 | 0.127 | 0.268 | NA |
| Post-heatwave | CORT release rate | Temperature | Gaussian (lmer) | 33.25 | 3 | **<0.001** | 0.667 | NA |
| 17°C | CORT release rate | Heatwave | Gaussian (lmer) | 1.6 | 1 | 0.205 | 0.243 | NA |
| 20°C | CORT release rate | Heatwave | Gaussian (lmer) | 1.49 | 1 | 0.222 | 0.1 | 0.473 |
| 23°C | CORT release rate | Heatwave | Gaussian (lmer) | 11.09 | 1 | **<0.001** | 0.479 | 0.572 |
| 26°C | CORT release rate | Heatwave | Gaussian (lmer) | 23.67 | 1 | **<0.001** | 0.703 | NA |
| Larvae (NF57) | Survival | Temperature | Binomial (glmer) | 1.67 | 4 | 0.796 | 0.07 | NA |
| Metamorph (NF66) | Survival | Temperature | Binomial (glmer) | 12.19 | 4 | **0.016** | 0.612 | 0.751 |
| Juvenile (150 dah) | Survival | Temperature | Binomial (Firth logistf) | 80.53 | 4 | **<0.001** | NA | NA |

**Table S2.** All significant Tukey-adjusted pairwise comparisons. Shown are estimated differences between temperature treatments and stages, standard errors (SE), degrees of freedom (df) are reported as estimated values, test statistics (t/z ratio), and p-values. Significant results (p < 0.05) indicate meaningful differences between temperature treatments within each trait and stage.

| Stage | Dependent variable | Contrast | Est | SE | Df | Test statistic | p-value |
| --- | --- | --- | --- | --- | --- | --- | --- |
| Larvae | Developmental rate (NF57-NF45)/dah | 17 - 20 | -0.154 | 0.750585 | 25 | 12.61238154 | **<0.001** |
|  |  | 17 - 23 | -0.271 | 0.750585 | 25 | 18.16360581 | **<0.001** |
|  |  | 17 - 26 | -0.391 | 0.750585 | 25 | 22.0716677 | **<0.001** |
|  |  | 17 - 29 | -0.375 | 0.750585 | 25 | 21.62756976 | **<0.001** |
|  |  | 20 - 23 | -0.117 | 0.750585 | 25 | 5.551224271 | **<0.001** |
|  |  | 20 - 26 | -0.237 | 0.750585 | 25 | 9.459286157 | **<0.001** |
|  |  | 20 - 29 | -0.221 | 0.750585 | 25 | 9.015188215 | **<0.001** |
|  |  | 23 - 26 | -0.12 | 0.750585 | 25 | 3.908061886 | **0.005175** |
|  |  | 23 - 29 | -0.104 | 0.750585 | 25 | 3.463963945 | **0.015111** |
|  | Body Mass | 17 - 23 | 376.3728 | 48.97929 | 24 | 7.68432564 | **<0.001** |
|  |  | 17 - 26 | 473.4221 | 45.59724 | 24 | 10.38269386 | **<0.001** |
|  |  | 17 - 29 | 471.8268 | 42.88602 | 24 | 11.00187726 | **<0.001** |
|  |  | 20 - 23 | 358.8673 | 43.4418 | 24 | 8.260875282 | **<0.001** |
|  |  | 20 - 26 | 455.9167 | 42.69915 | 24 | 10.67741678 | **<0.001** |
|  |  | 20 - 29 | 454.3213 | 47.14919 | 24 | 9.635823216 | **<0.001** |
|  | Q10 | 17 - 26 | -0.17028 | 0.041368 | 25 | -4.116209228 | **0.003092** |
|  |  | 17 - 29 | -0.31361 | 0.041368 | 25 | -7.581077028 | **<0.001** |
|  |  | 20 - 26 | -0.22444 | 0.041368 | 25 | -5.425606943 | **<0.001** |
|  |  | 20 - 29 | -0.36778 | 0.041368 | 25 | -8.890474743 | **<0.001** |
|  |  | 23 - 26 | -0.13278 | 0.041368 | 25 | -3.209703117 | **0.027255** |
|  |  | 23 - 29 | -0.27611 | 0.041368 | 25 | -6.674570917 | **<0.001** |
|  |  | 26 - 29 | -0.14333 | 0.041368 | 25 | -3.4648678 | **0.015079** |
|  | RMR | 17 - 23 | -0.00592 | 0.000971 | 25 | -6.097386796 | **<0.001** |
|  |  | 17 - 26 | -0.0083 | 0.000971 | 25 | -8.545493315 | **<0.001** |
|  |  | 17 - 29 | -0.01311 | 0.000971 | 25 | -13.49318523 | **<0.001** |
|  |  | 20 - 23 | -0.00367 | 0.000971 | 25 | -3.775117529 | **0.007165** |
|  |  | 20 - 26 | -0.00604 | 0.000971 | 25 | -6.223224047 | **<0.001** |
|  |  | 20 - 29 | -0.01085 | 0.000971 | 25 | -11.17091596 | **<0.001** |
|  |  | 23 - 29 | -0.00718 | 0.000971 | 25 | -7.395798431 | **<0.001** |
|  |  | 26 - 29 | -0.00481 | 0.000971 | 25 | -4.947691912 | **<0.001** |
|  | SVL | 17 - 23 | 2.361404 | 0.549025 | 24 | 4.301083412 | **0.002085** |
|  |  | 17 - 26 | 2.468713 | 0.511115 | 24 | 4.830057462 | **<0.001** |
|  |  | 17 - 29 | 3.028655 | 0.480724 | 24 | 6.300197598 | **<0.001** |
|  |  | 20 - 23 | 2.54269 | 0.486954 | 24 | 5.221625394 | **<0.001** |
|  |  | 20 - 26 | 2.65 | 0.478629 | 24 | 5.53664577 | **<0.001** |
|  |  | 20 - 29 | 3.209942 | 0.528511 | 24 | 6.073555587 | **<0.001** |
| Metamorph | Growth rate | 17 - 20 | -7.98147 | 2.098638 | 24 | -3.803165771 | **0.007015** |
|  |  | 17 - 23 | -11.4165 | 2.254299 | 24 | -5.064339773 | **<0.001** |
|  |  | 17 - 26 | -17.9432 | 2.098638 | 24 | -8.549904854 | **<0.001** |
|  |  | 17 - 29 | -9.38223 | 1.973853 | 24 | -4.753257447 | **<0.001** |
|  |  | 20 - 26 | -9.96169 | 1.965253 | 24 | -5.068910356 | **<0.001** |
|  |  | 23 - 26 | -6.52662 | 1.999433 | 24 | -3.264235378 | **0.024758** |
|  |  | 26 - 29 | 8.560926 | 2.170068 | 24 | 3.945003613 | **0.004982** |
|  | Body mass | 17 - 23 | 273.5234 | 90.67794 | 24 | 3.016427129 | **0.042944** |
|  |  | 17 - 26 | 250.8119 | 84.41657 | 24 | 2.971121694 | **0.047373** |
|  |  | 17 - 29 | 476.0331 | 79.39715 | 24 | 5.995594613 | **<0.001** |
|  |  | 20 - 29 | 287.1101 | 87.28978 | 24 | 3.289160906 | **0.023396** |
|  | SVL | 17 - 29 | 3.463938 | 0.605911 | 24 | 5.71690703 | **<0.001** |
|  |  | 20 - 29 | 3.264133 | 0.666143 | 24 | 4.900048445 | **<0.001** |
|  |  | 23 - 29 | 2.247563 | 0.721363 | 24 | 3.115717884 | **0.034534** |
|  |  | 26 - 29 | 2.04191 | 0.666143 | 24 | 3.065273662 | **0.038597** |
| All stages | CTmax | Juvenile 17 - Larvae 20 | 2.482035 | 0.595422 | 90.32711 | 4.168528471 | **0.005828** |
|  |  | Juvenile 17 - Metamorph 26 | -2.18375 | 0.614278 | 100.294 | -3.554991437 | **0.039928** |
|  |  | Juvenile 17 - Metamorph 29 | -2.40729 | 0.618214 | 102.1383 | -3.893935991 | **0.013782** |
|  |  | Juvenile 20 - Larvae 23 | 1.876796 | 0.470043 | 60.81685 | 3.992818988 | **0.013362** |
|  |  | Juvenile 23 - Larvae 26 | 1.667422 | 0.437543 | 47.97771 | 3.810872586 | **0.026505** |
|  |  | Larvae 17 - Juvenile 17 | -2.96965 | 0.595342 | 90.28209 | -4.988146934 | **<0.001** |
|  |  | Larvae 17 - Juvenile 20 | -4.3245 | 0.432784 | 47.09843 | -9.992268544 | **<0.001** |
|  |  | Larvae 17 - Juvenile 23 | -4.46689 | 0.404933 | 37.31592 | -11.03119833 | **<0.001** |
|  |  | Larvae 17 - Juvenile 26 | -4.95549 | 0.439083 | 49.33788 | -11.28600245 | **<0.001** |
|  |  | Larvae 17 - Juvenile 29 | -4.7904 | 0.787035 | 89.62844 | -6.086640246 | **<0.001** |
|  |  | Larvae 17 - Larvae 23 | -2.4477 | 0.36318 | 53.22295 | -6.739640727 | **<0.001** |
|  |  | Larvae 17 - Larvae 26 | -2.79947 | 0.368168 | 55.50749 | -7.603786751 | **<0.001** |
|  |  | Larvae 17 - Larvae 29 | -3.56959 | 0.372039 | 57.30863 | -9.594676766 | **<0.001** |
|  |  | Larvae 17 - Metamorph 20 | -3.54354 | 0.384188 | 67.02141 | -9.223451585 | **<0.001** |
|  |  | Larvae 17 - Metamorph 23 | -4.44851 | 0.383506 | 66.67657 | -11.59957912 | **<0.001** |
|  |  | Larvae 17 - Metamorph 26 | -5.1534 | 0.383547 | 66.6975 | -13.4361649 | **<0.001** |
|  |  | Larvae 17 - Metamorph 29 | -5.37694 | 0.387781 | 68.84603 | -13.86592959 | **<0.001** |
|  |  | Larvae 20 - Juvenile 20 | -3.83688 | 0.433958 | 47.5062 | -8.841600944 | **<0.001** |
|  |  | Larvae 20 - Juvenile 23 | -3.97927 | 0.405582 | 37.51429 | -9.81127032 | **<0.001** |
|  |  | Larvae 20 - Juvenile 26 | -4.46787 | 0.440243 | 49.74137 | -10.14865383 | **<0.001** |
|  |  | Larvae 20 - Juvenile 29 | -4.30278 | 0.787998 | 89.91232 | -5.460394011 | **<0.001** |
|  |  | Larvae 20 - Larvae 23 | -1.96008 | 0.362348 | 52.84612 | -5.40939443 | **<0.001** |
|  |  | Larvae 20 - Larvae 26 | -2.31185 | 0.367172 | 55.04809 | -6.296374139 | **<0.001** |
|  |  | Larvae 20 - Larvae 29 | -3.08198 | 0.370936 | 56.79292 | -8.308651105 | **<0.001** |
|  |  | Larvae 20 - Metamorph 20 | -3.05593 | 0.304733 | 265.856 | -10.02820953 | **<0.001** |
|  |  | Larvae 20 - Metamorph 23 | -3.96089 | 0.383571 | 66.70929 | -10.32635999 | **<0.001** |
|  |  | Larvae 20 - Metamorph 26 | -4.66579 | 0.383634 | 66.74114 | -12.16208745 | **<0.001** |
|  |  | Larvae 20 - Metamorph 29 | -4.88932 | 0.38728 | 68.59096 | -12.6247583 | **<0.001** |
|  |  | Larvae 23 - Juvenile 23 | -2.01919 | 0.430159 | 45.44341 | -4.694061386 | **0.002047** |
|  |  | Larvae 23 - Juvenile 26 | -2.50779 | 0.475932 | 62.89198 | -5.269208051 | **<0.001** |
|  |  | Larvae 23 - Metamorph 23 | -2.00081 | 0.317829 | 271.2998 | -6.29521766 | **<0.001** |
|  |  | Larvae 23 - Metamorph 26 | -2.7057 | 0.395457 | 72.79763 | -6.841965952 | **<0.001** |
|  |  | Larvae 23 - Metamorph 29 | -2.92924 | 0.384728 | 67.29459 | -7.613780953 | **<0.001** |
|  |  | Larvae 26 - Juvenile 26 | -2.15602 | 0.485354 | 66.57126 | -4.442154974 | **0.002912** |
|  |  | Larvae 26 - Metamorph 26 | -2.35393 | 0.324672 | 273.7588 | -7.250185795 | **<0.001** |
|  |  | Larvae 26 - Metamorph 29 | -2.57747 | 0.386677 | 68.2837 | -6.665689034 | **<0.001** |
|  |  | Larvae 29 - Metamorph 29 | -1.80734 | 0.309866 | 268.1134 | -5.832652649 | **<0.001** |
|  |  | Metamorph 17 - Juvenile 17 | -2.73741 | 0.616511 | 100.9448 | -4.440162481 | **0.002045** |
|  |  | Metamorph 17 - Juvenile 20 | -4.09225 | 0.442446 | 52.66932 | -9.249157897 | **<0.001** |
|  |  | Metamorph 17 - Juvenile 23 | -4.23465 | 0.42613 | 46.07442 | -9.937451098 | **<0.001** |
|  |  | Metamorph 17 - Juvenile 26 | -4.72324 | 0.448544 | 55.15578 | -10.53017494 | **<0.001** |
|  |  | Metamorph 17 - Juvenile 29 | -4.55815 | 0.786471 | 90.88189 | -5.79570397 | **<0.001** |
|  |  | Metamorph 17 - Larvae 23 | -2.21546 | 0.413051 | 82.0829 | -5.363640137 | **<0.001** |
|  |  | Metamorph 17 - Larvae 26 | -2.56722 | 0.420316 | 85.99097 | -6.107850944 | **<0.001** |
|  |  | Metamorph 17 - Larvae 29 | -3.33735 | 0.425631 | 88.87107 | -7.840953403 | **<0.001** |
|  |  | Metamorph 17 - Metamorph 20 | -3.3113 | 0.414458 | 87.59923 | -7.989458934 | **<0.001** |
|  |  | Metamorph 17 - Metamorph 23 | -4.21626 | 0.416644 | 88.80739 | -10.11957623 | **<0.001** |
|  |  | Metamorph 17 - Metamorph 26 | -4.92116 | 0.41631 | 88.62268 | -11.82089617 | **<0.001** |
|  |  | Metamorph 17 - Metamorph 29 | -5.14469 | 0.430183 | 96.33995 | -11.95931089 | **<0.001** |
|  |  | Metamorph 20 - Metamorph 26 | -1.60986 | 0.412707 | 86.63287 | -3.900741179 | **0.014526** |
|  |  | Metamorph 20 - Metamorph 29 | -1.83339 | 0.420327 | 90.84898 | -4.361826709 | **0.002944** |
|  |  | Metamorph 23 - Larvae 26 | 1.649037 | 0.399724 | 75.0227 | 4.125433024 | **0.007607** |
|  |  | Metamorph 26 - Larvae 29 | 1.58381 | 0.404214 | 77.38334 | 3.91824897 | **0.014552** |
|  | Life stage specific fat reserves | Larvae 17 - Larvae 23 | 0.004461 | 0.001261 | 60.36313 | 3.538017938 | **0.03431** |
|  |  | Larvae 17 - Larvae 26 | 0.00607 | 0.001388 | 75.04772 | 4.374092219 | **0.002155** |
|  |  | Larvae 17 - Metamorph 17 | -0.01927 | 0.001674 | 237.2442 | -11.51199043 | **<0.001** |
|  |  | Larvae 17 - Metamorph 23 | -0.00512 | 0.001439 | 94.07541 | -3.55509983 | **0.027723** |
|  |  | Larvae 20 - Metamorph 20 | -0.0062 | 0.001778 | 237.8335 | -3.485166765 | **0.028358** |
|  |  | Larvae 20 - Metamorph 23 | -0.00858 | 0.001671 | 107.6468 | -5.133331594 | **<0.001** |
|  |  | Larvae 20 - Metamorph 26 | -0.00628 | 0.001539 | 100.3693 | -4.082052955 | **0.004868** |
|  |  | Larvae 23 - Metamorph 23 | -0.00958 | 0.001525 | 227.4485 | -6.279838907 | **<0.001** |
|  |  | Larvae 23 - Metamorph 26 | -0.00728 | 0.001446 | 95.53312 | -5.035836434 | **<0.001** |
|  |  | Larvae 26 - Metamorph 26 | -0.00889 | 0.00156 | 226.2925 | -5.701108995 | **<0.001** |
|  |  | Metamorph 17 - Juvenile 17 | 0.019202 | 0.002549 | 145.6338 | 7.532651181 | **<0.001** |
|  |  | Metamorph 17 - Juvenile 20 | 0.018688 | 0.001613 | 66.45705 | 11.58884697 | **<0.001** |
|  |  | Metamorph 17 - Juvenile 23 | 0.018461 | 0.001581 | 60.64885 | 11.67407095 | **<0.001** |
|  |  | Metamorph 17 - Juvenile 26 | 0.019918 | 0.001656 | 74.38984 | 12.02671755 | **<0.001** |
|  |  | Metamorph 17 - Larvae 20 | 0.022727 | 0.002065 | 123.614 | 11.00679596 | **<0.001** |
|  |  | Metamorph 17 - Larvae 23 | 0.023729 | 0.00184 | 116.7298 | 12.89659494 | **<0.001** |
|  |  | Metamorph 17 - Larvae 26 | 0.025338 | 0.002048 | 133.0289 | 12.37469451 | **<0.001** |
|  |  | Metamorph 17 - Metamorph 20 | 0.016531 | 0.001633 | 128.7759 | 10.12390156 | **<0.001** |
|  |  | Metamorph 17 - Metamorph 23 | 0.014151 | 0.001691 | 130.7141 | 8.367515451 | **<0.001** |
|  |  | Metamorph 17 - Metamorph 26 | 0.016446 | 0.001811 | 134.2194 | 9.079038487 | **<0.001** |
|  |  | Metamorph 20 - Larvae 23 | 0.007198 | 0.001603 | 105.2842 | 4.490283588 | **0.001065** |
|  |  | Metamorph 20 - Larvae 26 | 0.008807 | 0.001776 | 121.7831 | 4.958660433 | **<0.001** |
|  |  | Metamorph 23 - Larvae 26 | 0.011187 | 0.001677 | 116.2895 | 6.669560201 | **<0.001** |
| 23°C | CORT release rate corrected | Baseline - Heatwave | -0.02281 | 0.00697 | 7.240258 | -3.273320391 | **0.012981** |
| 26°C | CORT release rate corrected | Baseline - Heatwave | -0.03576 | 0.007515 | 7.302598 | -4.758145374 | **0.001838** |
| Post-heatwave | CORT release rate corrected | 17 - 26 | -0.0386 | 0.008919 | 4.578213 | -4.327582073 | **0.032143** |
| Post-heatwave | CORT release rate corrected | 20 - 26 | -0.04584 | 0.009153 | 6.378886 | -5.008476268 | **0.008278** |

**Table S3.** Significant pairwise comparisons for survival to 150 days after hatching (dah), estimated using Firth penalized logistic regression. This endpoint showed complete separation (100% survival at 23°C) that prevented reliable estimation via standard mixed-effects logistic regression (see Methods); the random effect for Replicate was already estimated at zero variance in that model, so no random-effect information is lost by its omission here. Pairwise comparisons were computed as separate two-group Firth models for each pair of temperatures rather than derived from a single multi-level model, and p-values are therefore NOT Tukey-adjusted for multiple comparisons (compare Table S2, which is Tukey-adjusted). Estimates are log-odds differences between temperature treatments. Significant results (p < 0.05) are shown.

| Response variable | Contrast | estimate (log-odds) | p-value |
| --- | --- | --- | --- |
| Survival Juvenile 150 dah | 17 - 20 | -2.5905 | <0.001 |
|  | 17 - 23 | -3.439 | <0.001 |
|  | 17 - 26 | -2.1906 | <0.001 |
|  | 20 - 29 | 2.7867 | <0.001 |
|  | 23 - 29 | 3.6352 | <0.001 |
|  | 26 - 29 | 2.3868 | <0.001 |
